# Autorepression-based conditional gene expression system in yeast for variation-suppressed control of protein dosage

**DOI:** 10.1101/2022.12.05.519107

**Authors:** Aslı Azizoğlu, Cristina Loureiro, Jonathan Venetz, Roger Brent

## Abstract

Conditional control of gene expression allows an experimenter to investigate many aspects of a gene’s function. In the model organism *Saccharomyces cerevisiae*, a number of methods to control gene expression are widely practiced, including induction by metabolites, small molecules, and even light. However, all current methods suffer from at least one of a set of drawbacks, including need for specialized growth conditions, leaky expression, or the requirement of specialized equipment. Here we describe protocols using two transformations to construct strains that carry a new controller, in which all these drawbacks are overcome. In these strains, the expression of a controlled gene (gene of interest, or GOI) is repressed by the bacterial repressor TetR, and induced by anhydrotetracycline. TetR also regulates its own expression, creating an autorepression loop. This autorepression allows tight control of gene expression/ protein dosage with low cell to cell variation in expression. A second repressor, TetR-Tup1, prevents any leaky expression. We also present a protocol showing a particular workhorse application of such strains, to generate synchronized cell populations. We turn off the expression of the cell cycle regulator *CDC20* completely, arresting the cell population, and then back on so that the synchronized cells resume cell cycle progression. This control system can be applied to any endogenous or exogenous gene for precise expression.

**Basic Protocol 1:** Generating a parent WTC_846_ strain.

**Basic Protocol 2:** Generating a WTC_846_ strain with controlled expression of the targeted gene

**Alternate Protocol 1:** CRISPR-mediated promoter replacement

**Basic Protocol 3:** Cell cycle synchronization/Arrest and Release using the WTC_846-K3_::CDC20 strain

## INTRODUCTION

The conditional control of gene expression is a fundamental tool in biological research. It grants the researcher the ability to bring about the presence or absence of a gene product, or to tune its level of expression. Observing the resulting phenotypes frequently allows elucidation of biological function. The primary means used to effect conditional gene expression in eukaryotic cells like *Saccharomyces cerevisiae* are the use of promoters induced by chemical compounds, metabolites or other small molecules. These methods have have now been complemented by light-induced approaches. Transcription from chemically induced promoters depends on by native activator proteins (e.g. Gal4) or human-made activators (e.g. β-estradiol induced LexA-hER-B112) (Maya et al., 2008; Ottoz et al., 2014a; Taslimi et al., 2016; Xu et al., 2018). However these methods for conditional expression typically suffer from a number of drawbacks, including leaky expression (basal expression) in the absence of induction (Bellí et al., 1998a; Garí et al., 1997; Ottoz et al., 2014a; Taslimi et al., 2016; Xu et al., 2018), a reduction in cell doubling time caused by the activator (Gill & Ptashne, 1988), off-target induction of other genes (McIsaac et al., 2014a), and high levels of cell-to-cell variation of gene expression in isogenic populations (Elison et al., 2017a; Meurer et al., 2017a; Ottoz et al., 2014a; Xu et al., 2018). These limitations are particularly apparent when the research involves genes that are weakly expressed, assay of a null phenotype, or precise control of protein dosage to generate a well-behaved population to study phenotypes.

In previous work, we identified five desirable attributes for any conditional gene expression method for use in yeast, which, if made to be present together, would represent a marked improvement over previous approaches (Azizoğlu et al., 2021). These were (1) the ability to function in all commonly used *S. cerevisiae* growth media, (2) induction by an “orthogonal” small molecule that did not affect yeast cells, (3) lack of uninduced, basal expression (to allow generation of null phenotypes), (4) ability to generate a large range of precisely adjustable expression levels, and (5) have very high maximum expression to allow generation of overexpression phenotypes. To build this system, we modified an endogenous *S. cerevisiae* promoter to be repressed by the bacterial protein TetR, improved the repression efficiency by fusing TetR to an endogenous yeast repressor, and incorporated an autorepression loop in our system to reduce cell-to-cell variation. We recently described and characterized the resulting system in detail (Azizoğlu et al., 2021). We termed the collection of genetic elements that comprise this system a “well-tempered” controller, and named it WTC_846_, after Bach’s first Prelude and Fugue (C Major, BMV846) (Figure 1A).

**Figure 1.**
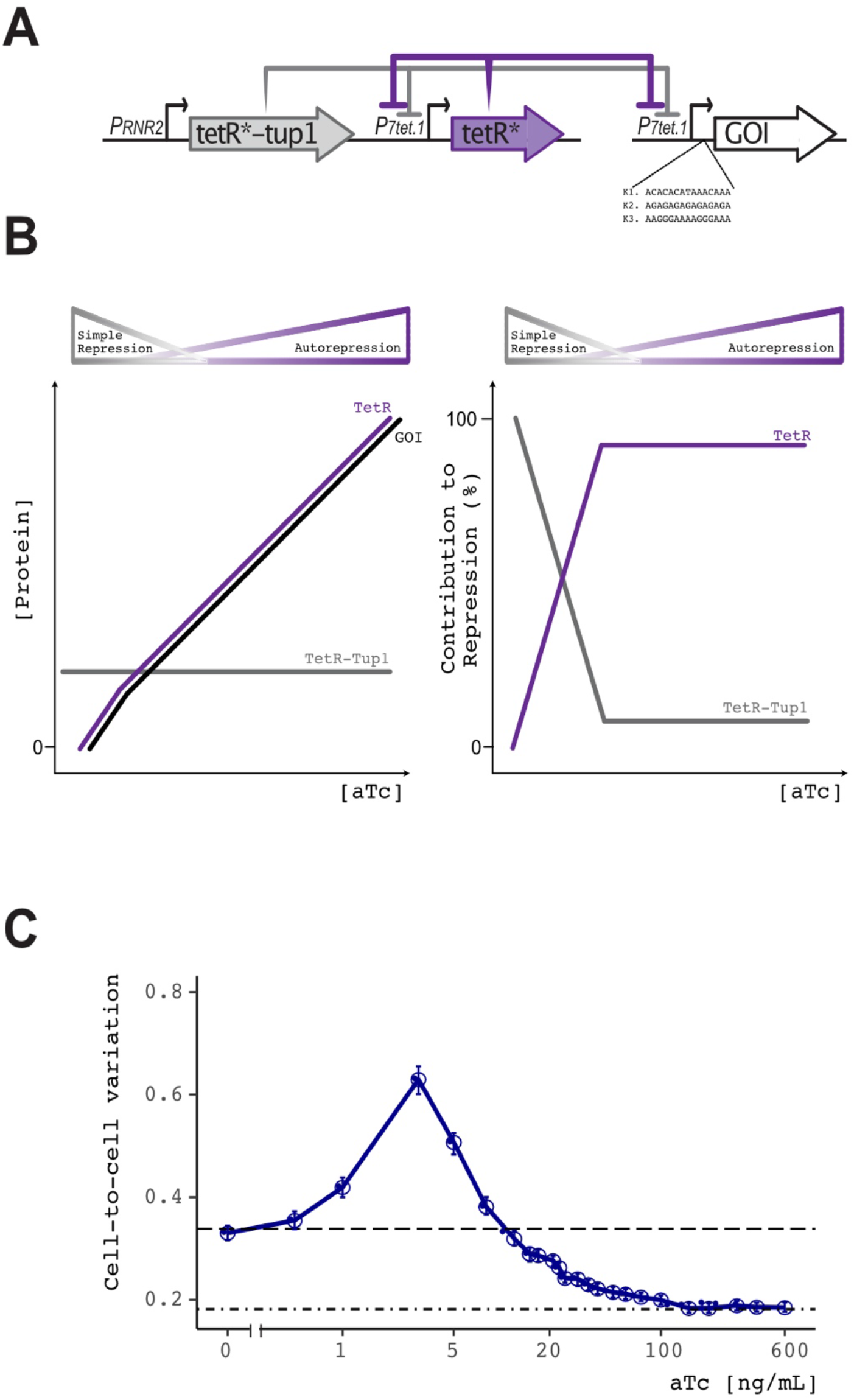
Architecture of WTC_846_ and its effects on cell-to-cell variation. **A) Complex Autorepression architecture of WTC_846_.** Schematic representation of the two repressors of WTC_846_ and their repressive effects on the different promoters of the system, shown as integrated constructs within the genome. Both TetR and TetR-Tup1 repress P_7tet.1_. However, only TetR and the Gene of Interest (GOI) are expressed from P_7tet.1_ and are subject to this repression. This creates an Autorepression loop, where TetR represses its own expression. The P_7tet_._1_ controlled gene and TetR are both expressed from instances of the same promoter and are, thus, in synchrony. TetR-Tup1 is not subject to Autorepression, since it is constitutively expressed from the RNR2 promoter. The three different Kozak sequences (last 15bp before the start codon) lead to different translation efficiencies and can be used to further fine tune the expression levels of the controlled gene. **B) Effect of aTc concentration on the expression of different elements of WTC_846_.** The left panel shows the protein concentrations inside the cell as aTc concentration increases. The higher the aTc concentration, the higher the levels of the autorepressed TetR and the protein encoded by the controlled gene (GOI). Their concentrations increase synchronously, since they are expressed by two instances of the same promoter, P_7tet_._1_. TetR-Tup1 is expressed constitutively; its concentration inside the cell is independent of aTc concentration. The right panel shows the contribution of each repressor to repression of P7_tet_._1_. At low aTc concentrations, TetR-Tup1 concentration is higher than TetR concentration. Therefore, TetR-Tup1 contributes more than TetR to repression of P_7tet_._1_. This is the Simple Repression regime of the dose response curve depicted above the plot. As aTc concentration increases, TetR contributes more and more to repression of P_7tet_ compared to TetR-Tup1, even as total repression of P_7tet_._1_ overall is decreasing. This is the Autorepression regime of the dose response curve. **C) Effect of aTc concentration on cell-to-cell variation in a strain where WTC_846_ controls Citrine expression**. The y axis indicates the variation in Citrine expression within a population measured by flow cytometry, after correcting for differences in cell size by fitting a linear model to the relationship between size and fluorescence as measured by flow cytometry (See (Azizoğlu et al., 2021) for a detailed explanation of cell-to-cell variation calculations). Variation peaks at low aTc concentrations when the Simple Repression architecture dominates. At higher concentrations of aTc, the Autorepression loop is active and suppresses this variation. The dashed line indicates the cell-to-cell variation of a strain where there is no Citrine expression. The dot-dash line indicates the variation in a strain where Citrine is expressed from the endogenous TDH3 promoter, and represents normal levels of variation in the wild type population. The error bars represent 95% confidence interval calculated using bootstrapping (n=1000). The experiment was repeated at least 3 times and a representative result is displayed. Figure was adapted with permission from (Azizoğlu et al., 2021).

The controller, WTC_846_ controls gene expression through two repressors, TetR and TetR-Tup1, which act on an artificial, TetR-repressed promoter, P_7tet.1_ (Figure 1). Derepression is achieved by addition of a tetracycline analog, anhydrotetracycline (aTc), to the medium. In WTC_846_ strains, P7tet.1 drives expression of a controlled gene (Gene of Interest or GOI), and a second instance of P_7tet.1_ controls expression of TetR itself. The consequence is that, at a given level of inducer, TetR and the controlled gene are expressed in synchrony (Figure 1B). This autorepression (AR) loop results in low cell-to-cell variation in expression of TetR and the controlled gene. However, the fact that controlled gene and TetR are both driven by instances of the same TetR-repressed promoter creates a problem, namely, that the controlled gene cannot be fully repressed. Therefore, WTC_846_ employs a second repressor that binds the same regulatory sites. This repressor, TetR-Tup1, is expressed constitutively outside of the AR loop, and so can fully repress P_7tet.1_. The strong repression afforded by TetR-Tup1 is likely due to the fact that the yeast Tup1 protein functions as powerful transcriptional repressor in yeast, operating by plural, reinforcing mechanisms of repression (Chen et al., 2013; Mennella et al., 2003; Zhang & Reese, 2004). The use of the two repressors, TetR and TetR-Tup1, allows complete repression of the controlled gene, but with the consequence that within the expression range between zero and the aTc concentration at which the AR loop activates, repression depends on TetR-Tup1 and cell-to-cell variation is higher (Figure 1C).

The system provides the researcher with an additional degree of control. WTC_846_ strains be constructed with one of 3 different extended Kozak sequences (the last 15bp before the start codon) for expression of the controlled gene, which confer varying translation efficiencies on the transcript of the controlled gene. Even though the WTC_846_ system has no detectable basal leakiness as assessed by flow cytometry or detectable by sensitive quantitative western blotting (Azizoğlu et al., 2021), certain genes that require only a few (<1000) copies of their protein to function, and also have long half-lives, might still be expressed enough to function at a baseline level. In such cases, using a low efficiency extended Kozak sequence to lower the translation efficiency has proved helpful to eliminate any such residual activity. On the other hand, for proteins that are very highly expressed in the cell, or for overexpression studies, high efficiency Kozak sequences can be used.

Here, we present a detailed protocol to allow researchers to construct strains in which WTC_846_ controls expression of a target gene in *S. cerevisiae* (Figures 2 & 3). Basic Protocol 1 describes the transformation required to incorporate the assembly of repressor elements into the genome of any auxotrophic *S. cerevisiae* starting strain. Basic Protocol 2 then explains how to replace the promoter of an endogenous target gene within the genome with the artificial promoter construct P_7tet.1_ using an antibiotic marker, such that the repressor elements incorporated in Basic Protocol 1 control the expression of this gene. We also present Alternate Protocol 1, which takes more time to carry out but is a transformation protocol with a higher success rate. Specifically, this alternative uses a Cas9-mediated method to increase promoter replacement efficiency, and allows replacement of the native promoter with WTC_846_ elements without introduction of an antibiotic resistance marker. Whether generated by Basic Protocol 2 or Alternate Protocol 1, expression of the targeted gene in any resulting WTC_846_ strain can then be precisely adjusted with the small molecule anhydrotetracycline, over a range of expression from complete absence to very high expression and with low cell-to-cell variation. We then describe Basic Protocol 3 as an example use case, in which we describe in detail the use of WTC_846_-regulated expression of the *CDC20* cell cycle progression gene to synchronize yeast cells in culture.

**Figure 2.**
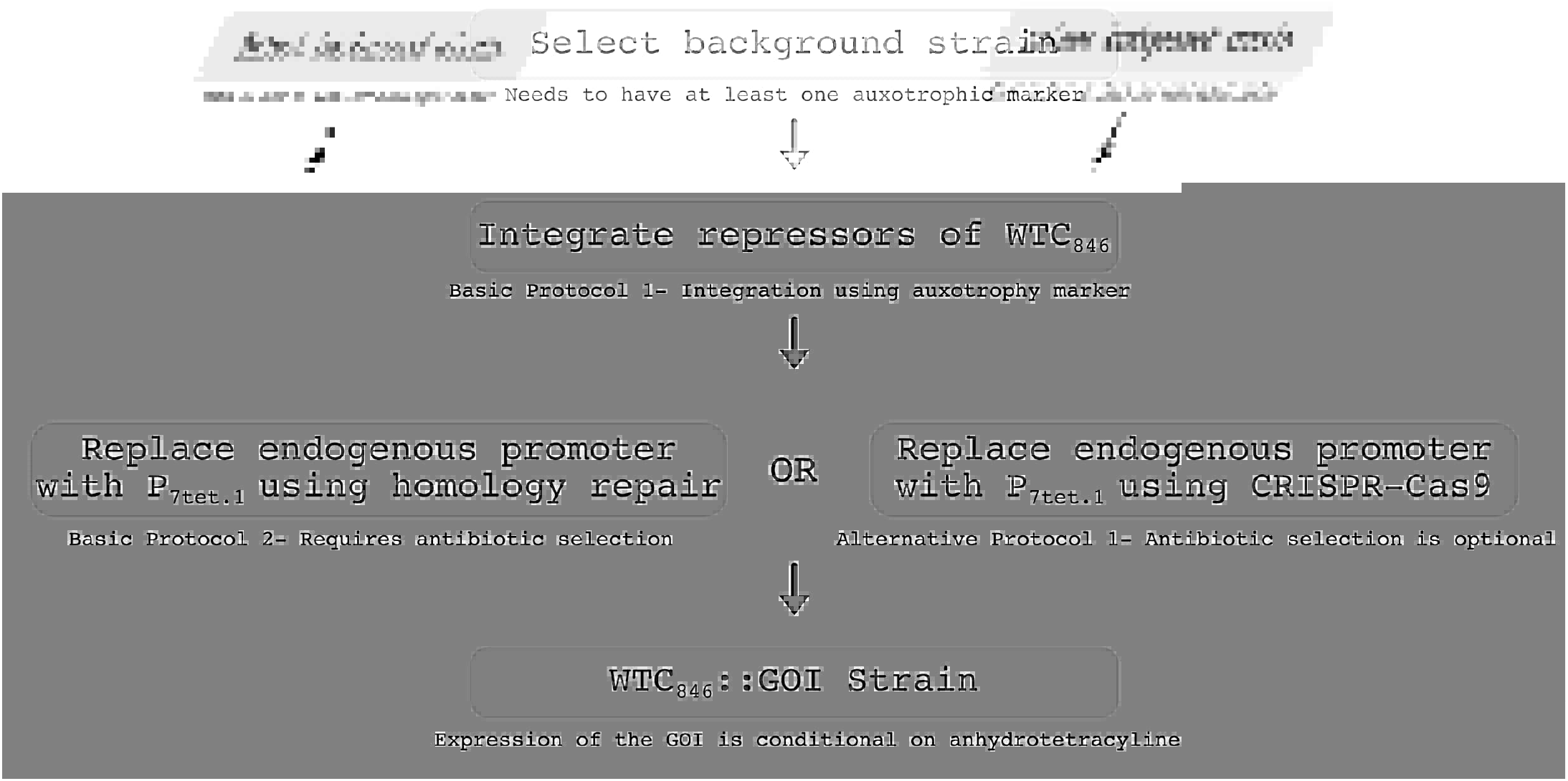
Workflow for generating WTC_846_ strains. A background strain is chosen with at least one auxotrophic marker deletion. The repressor elements of WTC_846_ found on a single integrative plasmid are integrated in the genome and the successful transformants are selected using the auxotrophic marker (Basic Protocol 1). That intermediate strain is then used to replace the promoter of an endogenous gene (Gene of Interest or GOI) with P_7tet.1_, using either a PCR template for homology repair (Basic Protocol 2), or a CRISPR-Cas9 method which increases efficiency of correct promoter replacement (Alternate Protocol 1). This results in the *WTC_846_::GOI* strain.

**Figure 3.**
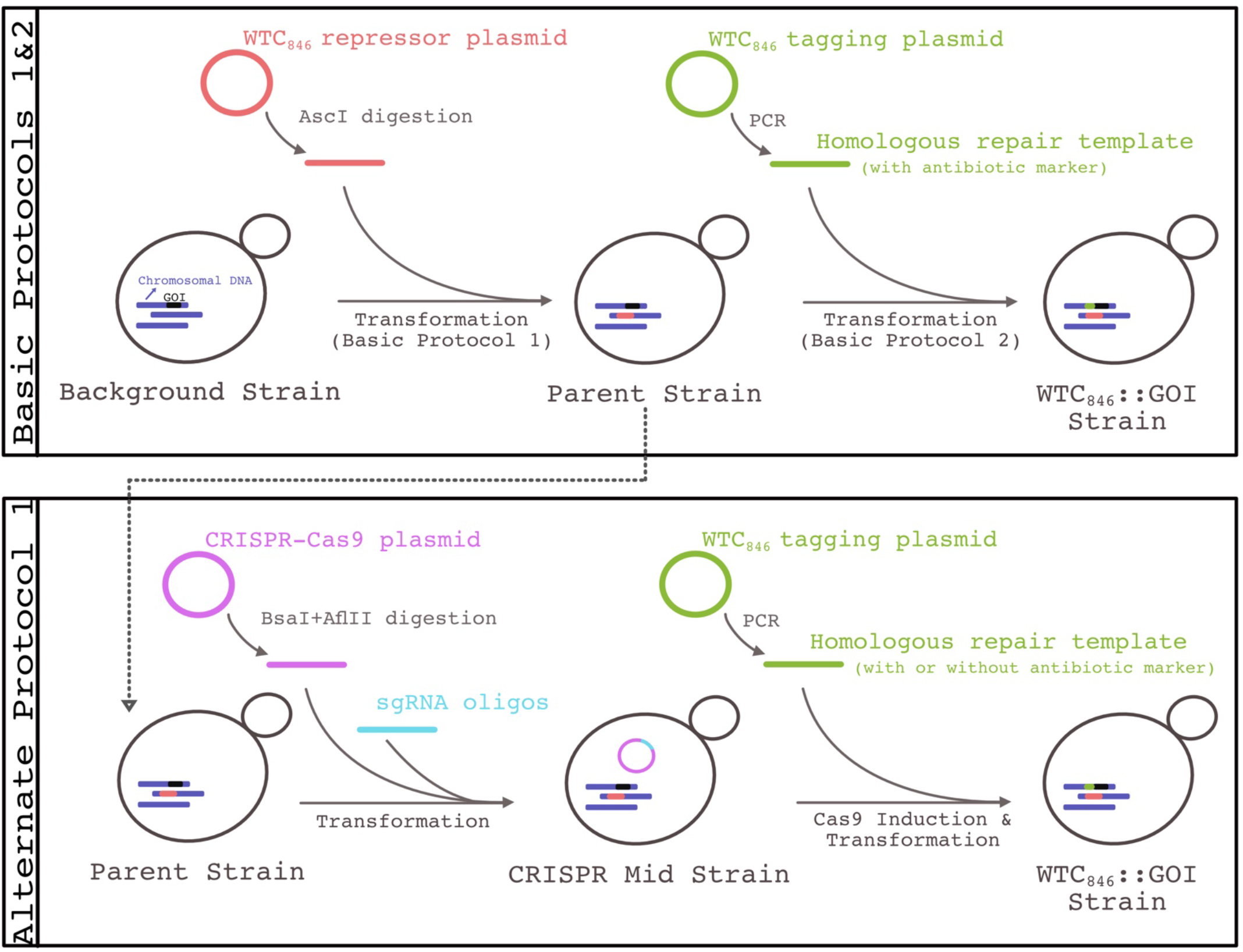
Overview of the protocols described. (Top) Basic Protocols 1&2 list the steps required to successfully place an endogenous gene of interest (GOI) under the control of WTC_846_. This involves two subsequent transformations, one to integrate the repressor elements (Basic Protocol 1) and a second one to replace the promoter of the endogenous GOI with P_7tet.1_ (Basic Protocol 2), to generate the strain where WTC_846_ controls the expression of the chosen gene (*WTC876::GOI*). (Bottom) Alternate Protocol 1 is a higher efficiency alternative to Basic Protocol 2. This involves two transformations. The first is to assemble a CRISPR-Cas9 plasmid together with guide RNAs that will then target the Cas9 enzyme to the promoter of the GOI. During the second transformation protocol, Cas9 expression is induced with β-estradiol and PCR product containing P_7tet_._1_ is introduced by transformation in order to replace the promoter of the GOI.

## STRATEGIC PLANNING

### Strain selection

The protocols described here have been tested with the widely used strains BY4741 and BY4742, which were derived from the progenitor strain S288C. S288C is a commonly used, non-flocculent yeast strain that is the ancestor of many strains used in research (Mortimer & Johnston, 1986), and its genome was the first eukaryotic genome to be fully sequenced (Goffeau et al., 1996). However, any *S.cerevisiae* strain with at least one of the auxotrophic marker deletions Δleu2, Δhis3, Δlys2, Δmet15 or Δura3 can be used to perform the protocols described below. Users can obtain strains from ATCC (atcc.org) or EUROSCARF (euroscarf.de).

### Determining Required Inducer Concentration

The end product of these protocols is a strain where the expression level of a the controlled gene (Gene of Interest, GOI) depends on the extracellular concentration of anhydrotetracycline (aTc) present in the medium. If the GOI encodes a protein native to *S. cerevisiae* and essential for growth, it will likely need to be expressed at endogenous levels when creating and maintaining this strain. Here we describe how to determine and appropriate aTc concentration for endogenous GOI in both liquid and solid media.

In the final strain, the GOI expression is controlled by the promoter P_7tet.1_. This promoter is based on the endogenous P_TDH3_ promoter of *S. cerevisiae* and has a similar maximum expression level, as assessed by fluorescence in strains where the promoters are drive expression of the a mCitrine reporter (Azizoğlu et al., 2021). Therefore, the maximum protein concentration reached by fully inducing WTC_846_ with 600 ng/mL aTc is roughly equivalent to *TDH3* protein concentration inside the cell, which is measured to be around 2 million in rich media (YPD) (Ho et al., 2018). Assuming that the GOI is under similar post-transcriptional regulation (e.g. degradation) as *TDH3*, protein levels will be approximately around 2 million molecules per cell at maximum induction of P_7tet.1_ in rich medium.

However, the actual expression level of the GOI will depend also on any potential translational and post-translational regulation that might exist. For example, if the degradation rate of the controlled protein is much faster than the approximately 12h half-life measured for *TDH3*(Christiano et al., 2014), the final controlled protein expression level is likely to be lower than 2 million molecules per cell at full induction.

Therefore, we suggest consulting the *Saccharomyces* Genome Database (SGD) (www.yeastgenome.org), where the endogenous expression levels and half-life information of many proteins are available under different media conditions (Cherry et al., 2012; Hellerstedt et al., 2017). Based on this information, an approximation can be made as to what percentage of maximal P_7tet.1_ induction is required to maintain endogenous levels of the GOI. Then, Figure 5 in conjunction with Table 3 can be used to estimate the required aTc concentration in liquid YPD media and Figure 6 for solid YPD and synthetic media to achieve this activation level. In general, synthetic media-based conditions require less aTc than rich media conditions to reach the same expression level (Azizoğlu et al., 2021) It is therefore recommended to apply a 5-10 fold reduction in aTc concentrations in synthetic liquid media compared to rich liquid media.

**Figure 4.**
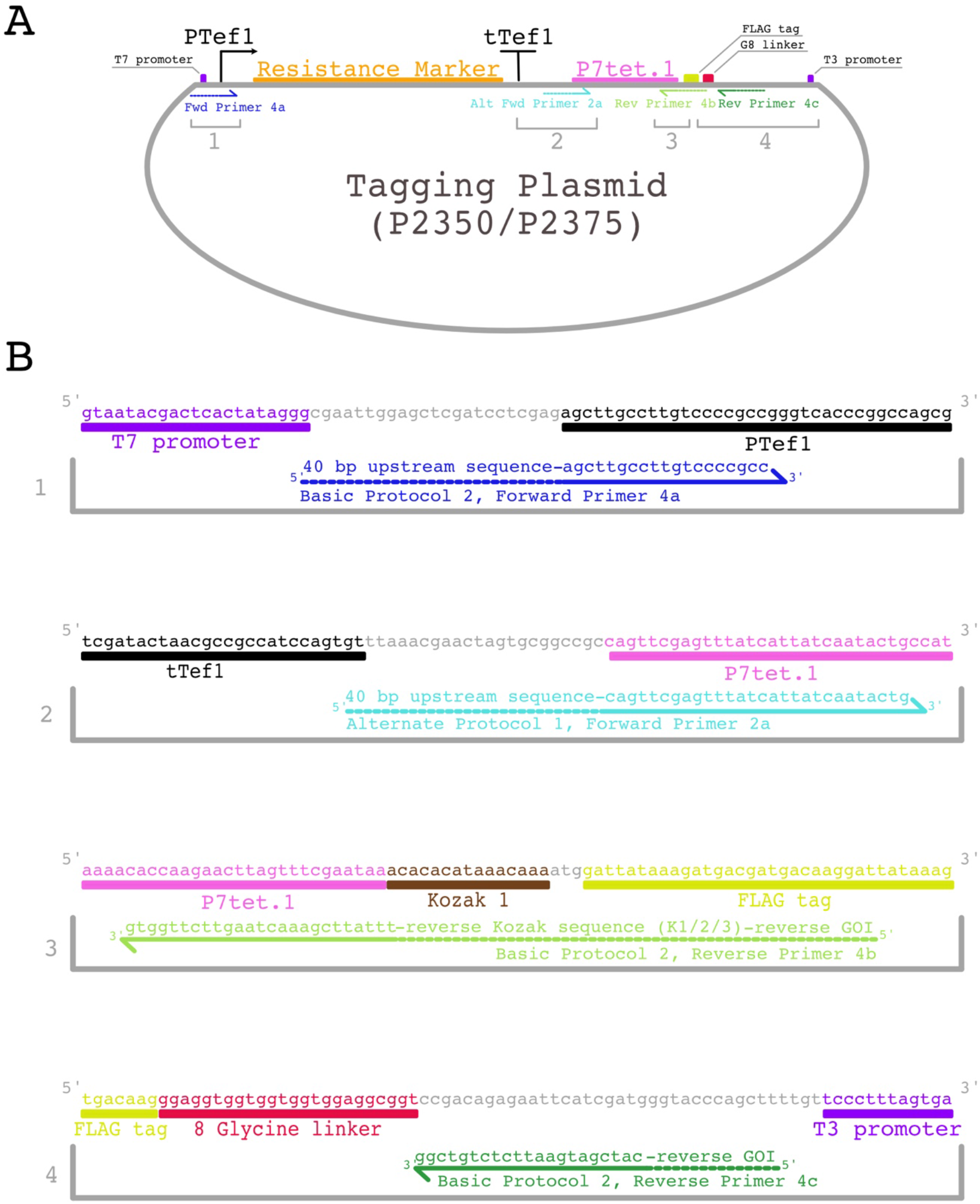
Overview of primer design for generating the P_7tet.1_tagging PCR product. **A) Schematic representation of the P_7tet.1_.-tagging plasmids P2350 and P2375**, showing the location of the resistance marker expression cassette, the P_7tet.1_. promoter, and the annealing positions of the different primers (not to scale). The primers are named either Forward (Fwd) or Reverse (Rev) to indicate direction, and are numbered after the protocol step in which they are described. The portions of the primers depicted with solid lines indicate annealing to the plasmid, while the dashed lines indicate tails. The brackets mark the regions that are depicted in more detail in (B). **B) The four regions where the different primers anneal, depicted in sequence-level detail.** The primer names refer to the protocol and the protocol step in which they are described, while Forward or Reverse indicate their direction. 5’ and 3’ ends of each sequence and each primer are labelled.

**Figure 5.**
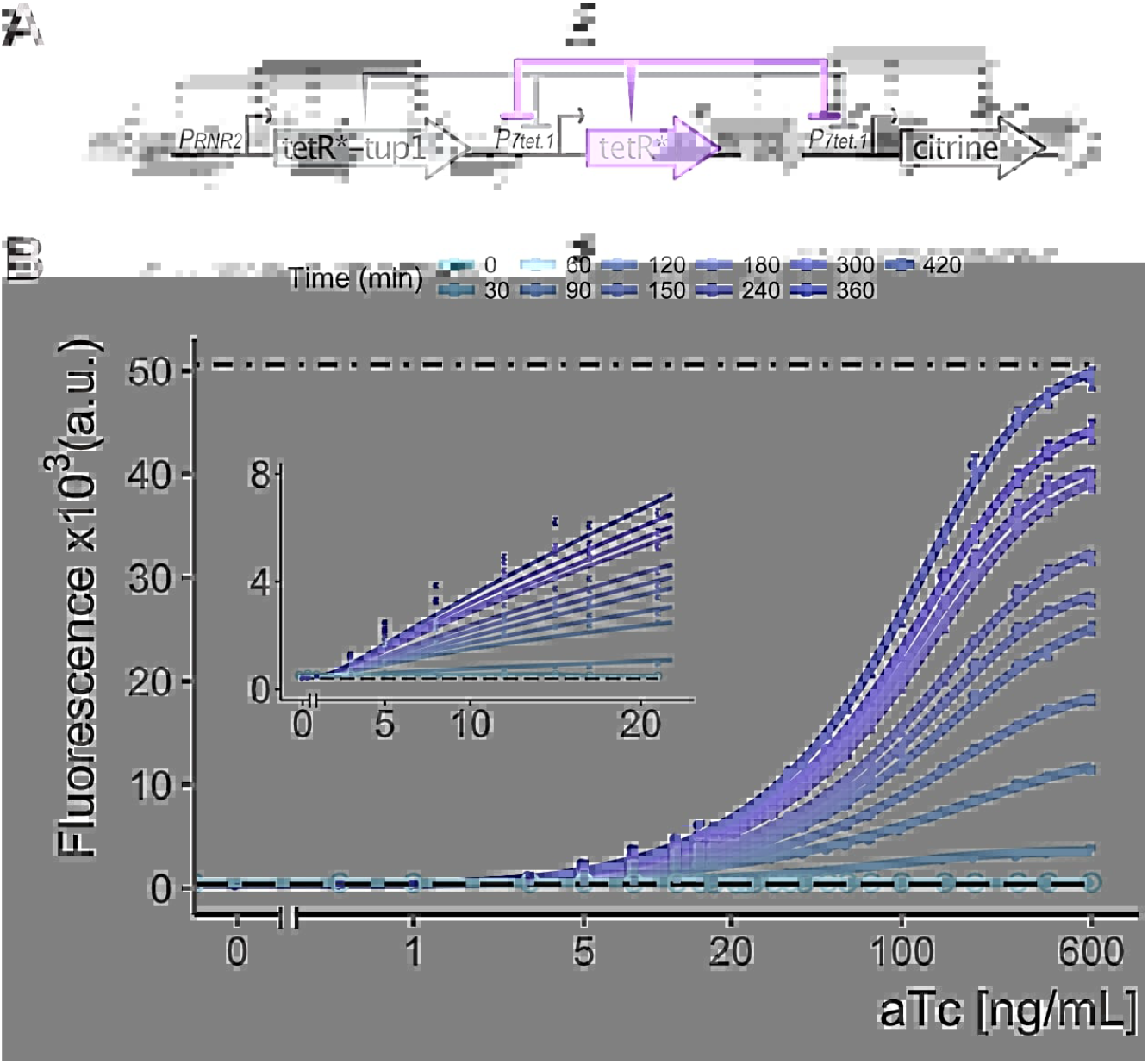
Time-dependent dose-response of the *WTC_846_::Citrine* strain. **A) Architecture of the *WTC_846_::Citrine* strain**. The repressor elements are comprised of a P_7tet.1_ -controlled TetR and a PAct1-controlled TetR-Tup1. (*) denotes a nuclear localization signal, and blunt ended arrows indicate repression. The repressor elements were integrated at the *URA3* locus. Citrine was placed under control of P_7tet.1_ and integrated at the *LEU2* locus.**B) Time-dependent dose response to anhydrotetracycline.** Citrine production was induced with different concentrations of anhydrotetracycline and Citrine fluorescence was measured using flow cytometry (488nm excitation, 530/30nm emission) every 30 minutes. Circles represent the median of the population and error bars indicate 95% confidence interval. The dot-dashed line indicates the fluorescence measured in a strain with only P_7tet.1_ controlled citrine integrated at the *LEU2* locus. The dashed line indicates the autofluorescence of a strain where no elements of the WTC_846_ was integrated. Numerical data is provided in Table 3 for time point 420 minutes. Figure was reproduced with permission from (Azizoğlu et al., 2021).

**Table 1.** Troubleshooting Guide for Yeast Transformation

| Problem | Possible Cause | Solution | Relevant Protocol |
| --- | --- | --- | --- |
| No colonies observed on the selection plate | Wrong dropout or antibiotic plate used. | Check that the auxotrophic/antibiotic marker on the transformed DNA and the dropout/antibiotic plate match correctly. | Basic Protocol 1 & 2, Alternative Protocol 1 |
|  | Too few cells were added to the transformation mix. | Make sure that the liquid cell culture has at least 10 million cells per mL at the start of the protocol or had reached an OD<sub>600</sub> of 2.0. | Basic Protocol 1 & 2, Alternative Protocol 1 |
|  | Cells were not in exponential phase at the start of the protocol. | Make sure that the cells have been growing in rich media for at least 3 hours and are still in exponential phase at the start of the protocol. This can be confirmed by checking the cells under light microscope, 20X or 40X magnification. Almost all the cells in the field of view should be budding. | Basic Protocol 1 & 2, Alternative Protocol 1 |
|  | Too little DNA transformed. | Increase the amount of DNA that goes into the transformation mix. This can apply to digested plasmids, PCR products, and double stranded oligos. | Basic Protocol 1 & 2, Alternative Protocol 1 |
| Too many colonies observed on the selection plate. | Wrong dropout or antibiotic plate used. | Check that the auxotrophic/antibiotic marker on the transformed DNA and the dropout/antibiotic plate match correctly. | Basic Protocol 1 & 2, Alternative Protocol 1 |
|  | Too high transformation efficiency, too many cells plated. | Repeat the protocol and try plating half or a quarter of the transformation mix, depending on how dense the plate was. | Basic Protocol 1 & 2, Alternative Protocol 1 |
| No correct colony can be found with colony PCR. | Incorrect oligos used. | If checking for plasmid integration, make sure the correct oligos are used. If checking for promoter replacement, make sure reverse oligo anneals at the start of the GOI, and doesn't have additional binding sites within the genome using the BLAST function on Saccharomyces Genome Database (yeastgenome.org). | Basic Protocol 1 & 2, Alternative Protocol 1 |
|  | Promoter replacement was unsuccessful. | If promoter replacement was performed with Basic Protocol 2, try Alternate Protocol 1 instead.<br>If Alternate Protocol 1 was used, make sure the CRISPR-Cas9 plasmid has correctly assembled by repeating the spotting assay and observing a clear difference in growth rates with and without β-estradiol.<br>Make sure the anhydrotetracycline concentration used in the selection plate is appropriate for the endogenous GOI expression level. Try higher and lower concentrations as necessary. | Basic Protocol 2 and Alternate Protocol 1 |
| Spotting assay shows no difference in growth rate between +/- β-estradiol plates | G418 was omitted from the plates. | Make sure G418 is present at the correct concentration on the spotting assay plates, otherwise the cells will have no selective pressure to maintain the CRISPR-Cas9 plasmid.<br>Do not use Kanamycin as an alternative to G418. Kanamycin does not enter *S. cerevisiae* cells. | Alternative Protocol 1 |
| Arrest is not complete. | Incomplete wash steps. | Increase the number of wash steps with media without inducer, and remove as much of the old media as possible after centrifugation.<br>If possible, start with a lower concentration of inducer. | Basic Protocol 3 |
|  | Too much protein product, too low degradation rate, or too low growth rate. | If possible, start with a lower concentration of inducer before inducing arrest.<br>If the growth rate is very slow due to the media conditions or the temperature of incubation, or if the protein product is very stable, arrest will take a long time. Wait longer and dilute as necessary. | Basic Protocol 3 |
| Induction is not synchronous. | Arrest wasn't complete. | See points above. | Basic Protocol 3 |
|  | Inducer concentration is too low. | If inducer concentration is very low, cells might initially take up unequal amounts. Try using a higher concentration of inducer for the release step. | Basic Protocol 3 |

**Table 2.**
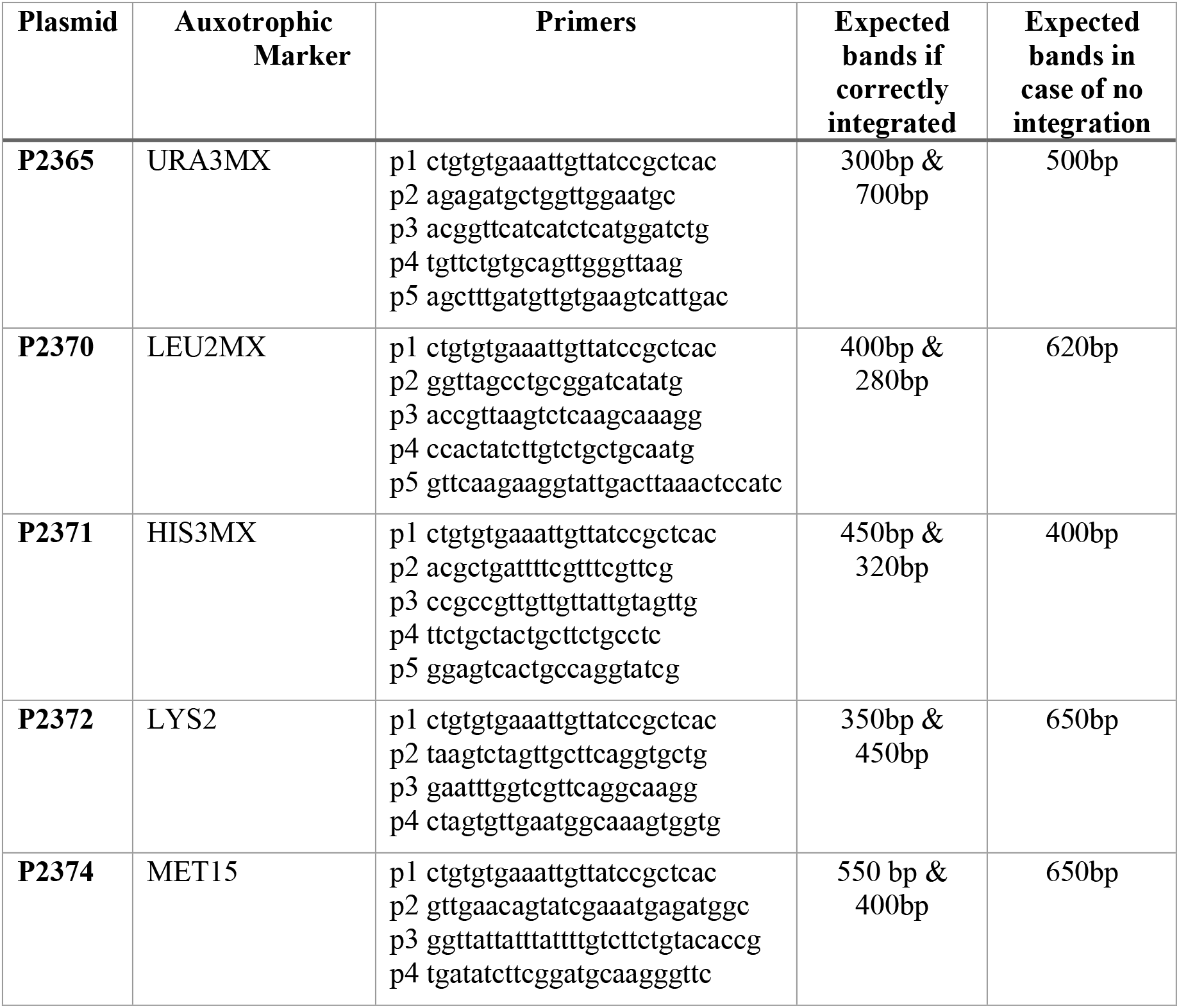
Colony primers to check for correct chromosomal integration, and the expected length of the PCR products. The repressor plasmids are based on the pRG shuttle vector series and the colony PCR primers and protocol are as described for this vector series in the original publication (Gnügge et al., 2016).

**Table 3.** Time dependent dose response of the *WTC_846_::Citrine* strain. Numerical data for the time point 420 minutes presented in Figure 5 Panel B.

| Dose [ng/mL] | Lower Confidence Interval | Upper Confidence Interval | Time (min) | Median |
| --- | --- | --- | --- | --- |
| 0 | 374 | 386 | 420 | 380 |
| 0.5 | 386 | 406 | 420 | 398 |
| 1 | 453.9875 | 478 | 420 | 464.5 |
| 3 | 1274.5 | 1345 | 420 | 1306.5 |
| 5 | 2439 | 2542.5 | 420 | 2483.5 |
| 8 | 3762.975 | 3933.5 | 420 | 3860.5 |
| 12 | 4818.9875 | 5004.5 | 420 | 4902.5 |
| 15 | 6098.5 | 6341.5 | 420 | 6214.5 |
| 17 | 5967.4875 | 6173 | 420 | 6081.5 |
| 21 | 6426 | 6684.025 | 420 | 6560.5 |
| 23 | 7311 | 7527.075 | 420 | 7457 |
| 25 | 7981.925 | 8309.575 | 420 | 8167 |
| 30 | 9277.9 | 9652.6375 | 420 | 9490 |
| 35 | 11147.95 | 11593 | 420 | 11366.5 |
| 40 | 11751.4875 | 12171.025 | 420 | 11957.5 |
| 50 | 14583.2625 | 15075.0625 | 420 | 14823.5 |
| 60 | 17076.5 | 17734.0625 | 420 | 17374.5 |
| 75 | 20527.925 | 21282.2875 | 420 | 20851 |
| 100 | 25402.8 | 26314 | 420 | 25866 |
| 150 | 33428.925 | 34546.8375 | 420 | 33982.5 |
| 200 | 40186.9375 | 41865 | 420 | 41008 |
| 300 | 44648.5 | 46346.7125 | 420 | 45470 |
| 400 | 46501.225 | 48226.5375 | 420 | 47350 |
| 600 | 48225.4875 | 50243.5 | 420 | 49338.5 |

**Figure 6.**
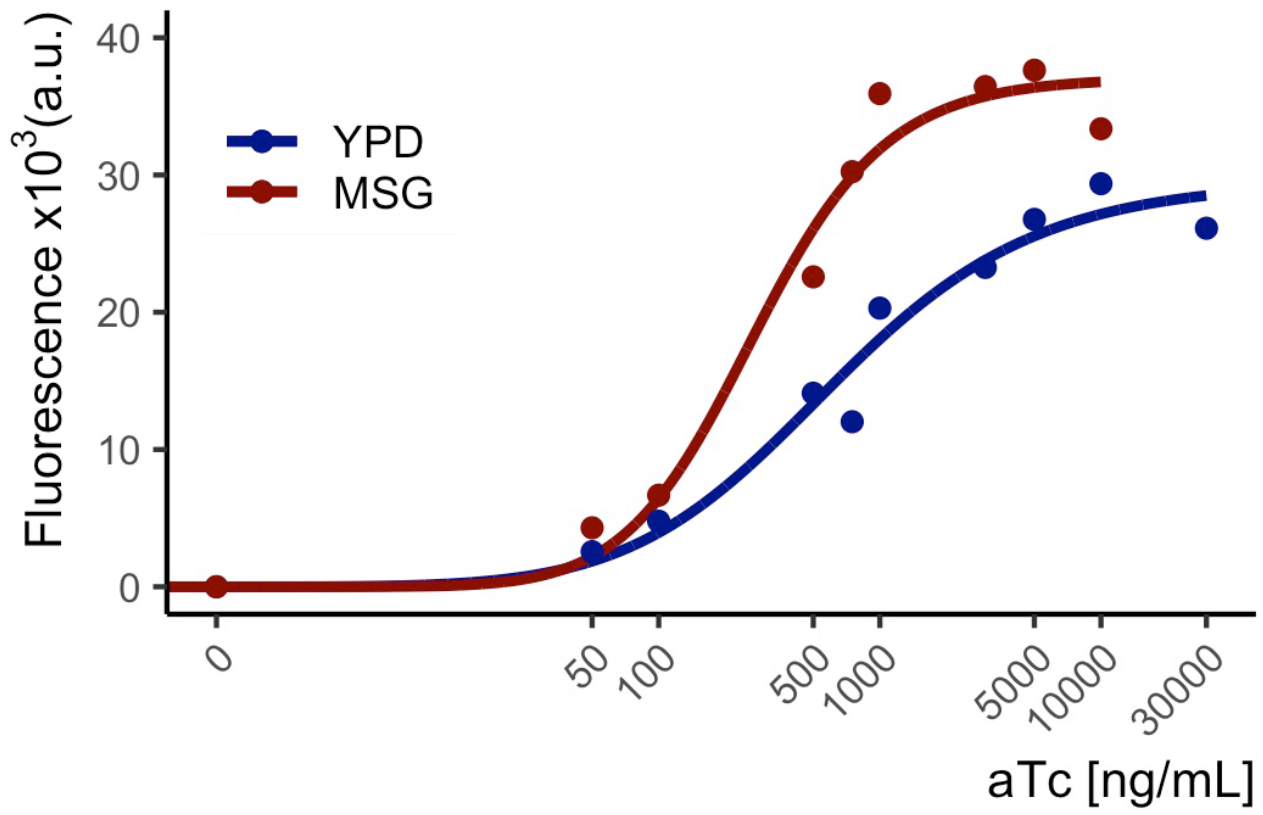
Dose response of *WTC_846_::Citrine* strain to anhydrotetracycline on solid media. Cells were streaked onto YPD or MSGfull plates containing different concentrations of aTc and grown for 2 days. Single colonies were resuspended in PBS and fluorescence was measured using flow cytometry (488nm excitation, 530/30nm emission). Circles represent the median of the population. MSGfull is a synthetic medium that contains 1% glutamine as the main nitrogen source, 2% glucose as the main carbon source, 1.7% Yeast Nitrogen Base, and a complete complement of essential and non-essential amino acids.

For example, if a protein has reported expression levels are around 200000 molecules per cell in rich media, this would be around 10% of *TDH3* levels. Based on fluorescence data shown in Table 3, this would correspond to an aTc dose of 12 ng/mL in liquid rich media, and around 1 to 2.5 ng/mL in liquid synthetic media. Based on Figure 6, solid rich media concentration would correspond to around 100ng/mL, whereas synthetic solid media concentration would be around half at 50ng/mL.

Most endogenous genes tested in our hands tolerated slight changes in their concentration well, therefore not necessitating a more precise estimation of aTc concentrations that would yield exact endogenous expression levels. Nevertheless, since the calculated aTc concentration will be an approximation, we recommend testing a range of concentrations from half the calculated amount to double the calculated amount. In cases where increased or decreased expression of the GOI results in an observable phenotype (e.g. changes in growth rate), this can be checked to determine the ideal concentration. Additionally, low expression levels measured by fluorescence is always obscured by autofluorescence, and therefore Table 3 cannot be used to estimate concentrations of very low expressed genes with the calculation method explained above. This applies to genes that are endogenously expressed at a level less than around 0.2% of *TDH3* levels. For such genes, we recommend testing 3 or 4 aTc concentrations ranging from 1-10 ng/mL for rich liquid media.

## BASIC PROTOCOL 1

### Generating a parent WTC_846_ strain

This multi-day protocol explains how to create a strain in which a single copy of the assembled repressor elements of the WTC_846_ system (Azizoğlu et al., 2021) is incorporated into the genome. Here, the user will decide on the auxotrophic marker to use, select the appropriate plasmid and order it from Addgene ahead of time. Plasmid preparation takes 3 days after it arrives. This protocol also requires yeast strain preparation steps three days in advance (Day −2) and one day in advance (Day 0) before the transformation can be performed. On the day the protocol is executed (Day 1), the chosen plasmid is transformed into the desired. *S. cerevisiae* starting strain background using an optimized LiAc-based transformation method. On Day 3, the colonies that appear are screened for correct integration using colony PCR. The resulting strain, hereafter called the **parent strain**, will then be used in subsequent protocols to generate strains in which the expression of the targeted gene is controlled by WTC_846_.

### Materials

*S. cerevisiae* cells (S288C or any other desired background)

YPD media (See Reagents and Solutions)

DNA Miniprep kit (ZymoPure, Zymo Research or similar)

AscI restriction enzyme (NEB, R0558S)

0.2M Lithium Acetate (CAS number 6108-17-4) solution, dissolved in ddH2O

4M Lithium Acetate solution, dissolved in ddH_2_O

0.4M Tris-HCl buffer (See Reagents and Solutions)

0.08M EDTA solution, dissolved in ddH_2_O.

Salmon Sperm DNA solution (See Reagents and Solutions)

Sterile ddH_2_O

0.2M Lithium Acetate + 20% (v/v) glycerol (CAS number 56-81-5) solution, dissolved in ddH_2_O

58% (w/v) PEG solution, dissolved in ddH2O (Average molecular weight 3350, CAS number 25322-68-3)

DMSO (CAS number 67-68-5)

TE buffer (0.1M Tris HCl Buffer, 0.01M EDTA solution)

Synthetic Defined Media (See Reagents and Solutions)

Synthetic Defined Media Dropout Plate (See Reagents and Solutions)

YPD plate (See Reagents and Solutions)

LB media with Carbenicillin (See Reagents and Solutions)

LB agar plate with Carbenicillin (See Reagents and Solutions)

70% (v/v) glycerol (CAS number 56-81-5) solution, dissolved in ddH_2_O

20mM NaOH (CAS number 1310-73-2) solution, dissolved in ddH_2_O.

5M Betaine (CAS number: 107-43-7) solution

Thermophol Buffer (NEB B9004)

2mM dNTP solution (Sigma, DNTP10)

Primers (See Table 2), IDT or similar

2% electrophoresis gel (See Reagents and Solutions)

DNA loading buffer (See Reagents and Solutions)

DNA Molecular Weight Marker (NEB 1kb Plus DNA ladder, cat. No. N3200, or equivalent)

TBE buffer (See Reagents and Solutions)

Taq DNA polymerase (NEB M0273)

Plasmids from Addgene encoding WTC_846_ repressor elements (See Step 1)

Cell counter (Beckman, Z2 Coulter or equivalent, optional)

OD_600_ measurement device (PerkinElmer, Lambda Bio or equivalent, as an alternative to the Cell Counter)

Centrifuge (Thermo Scientific, Hereaus Multifuge 3SR+ or equivalent)

50-mL Falcon tubes

100-mm Petri dishes

Glass culture tubes (55mL, Huberlab 9.6131.33 or equivalent)

Eppendorf Tubes (1.5 or 2mL)

Tabletop centrifuge (Eppendorf 5514R or equivalent)

Incubator (Binder Series B Classic Line or equivalent)

Shaking incubator (Kuhner Shaker ISF1-X or equivalent)

Vortex (Scientific Industries, Vortex Genie 2 or equivalent)

Rotating wheel (optional)

Thermomixer (Eppendorf Thermomixer Comfort or equivalent with heating and shaking functions)

PCR tubes (0.2mL volume)

Thermocycler

Electrophoresis gel chamber and power unit

**−80°C freezer**

### Protocol steps

1. Select an appropriate auxotrophic marker to use depending on the background of the chosen *S. cerevisiae* strain. Acquire the plasmid encoding WTC_846_ repressors corresponding to this marker well ahead of time. There are 5 versions of the plasmid, available from Addgene (https://www.addgene.org/browse/article/28203500/) with the following markers: URA3(P2365), LEU2(P2370), HIS3(P2371), LYS2(P2372), MET15(P2374). All plasmids are “shuttle vectors”, meaning they can be maintained in E. coli when grown in the presence of Carbenicillin or Ampicillin.
2. Streak the bacterial stab received from Addgene on an LB agar plate containing 100μg/mL Carbenicillin and incubate at 37 °C overnight.
3. Pick a single colony and grow in 4mL LB media with Carbenicillin overnight at 37°C.
4. Mix 1mL of the overnight culture with 0.5mL 70% glycerol to store at −80°C for future use. Inoculate a small amount of the stock into LB media with Carbenicillin whenever needed.
5. Miniprep and resuspend the selected repressor plasmid in double distilled, autoclaved water. This protocol requires 1μg of purified plasmid. *It is advisable to send the resulting purified plasmid for Sanger sequencing at a sequencing facility. The insert on the plasmid is flanked by the standard primer binding sites T7(*TAATACGACTCACTATAGGG*) and T3(*TTAACCCTCACTAAAGG*). Inspect the sequencing results to confirm that it matches the sequence indicated on Addgene.*

### Day −2 to Day 0: Strain Preparation for Yeast Transformation

**6. Day −2:** Streak the yeast strain to be transformed onto a YPD plate and incubate at 30°C for 2 days.
7. **Day 0:** Start a liquid culture of the yeast strain to be transformed by inoculating a single colony in 5 mL of rich media (YPD) in a glass culture tube. Shake overnight at 30°C, 275 rpm.
8. **(Day 0)** Digest about ~1 μg of the selected plasmid in a volume of 20 μL with the restriction enzyme AscI, following the enzyme’s manufacturer instructions. Incubate overnight at 37°C. Restriction digestion can also be performed on the day of the transformation (Day 1). However, the reaction should be incubated at 37°C for at least 2 hours.

### Day 1: Transformation

9. Dilute the overnight culture into 10 mL of YPD in a glass culture tube. If no cell counter is available, dilute down to an OD_600_ of 0.5 or just use a 1:50 dilution ratio (i.e. add 200 μL to the 10-mL culture). If one is available, dilute the culture to a concentration of 2.5 million cells/mL. Shake at 30°C for 3 hours at 275rpm until the culture reaches a concentration of 10 million cells/mL or an OD_600_ of 2.0.
10. Transfer the cells to a 50mL Falcon tube. Harvest the cells by centrifuging at 3000g for 3 minutes at room temperature. Remove supernatant without disrupting the cell pellet.
Resuspend pellet in 1 mL of 0.2M LiAc solution by pipetting or briefly vortexing, and centrifuge at 1000g for 1 minute at room temperature. Remove supernatant carefully.
11. During the centrifugation, prepare the transformation mix by combining the following sterile reagents in a 1.5 or 2mL Eppendorf tube (volumes given are for a one transformation reaction):

a. 20μL 4M LiAc solution
b. 10μL 0.4M Tris
c. 10μL 0.08M EDTA
d. 10μL ssDNA
e. 50μ L dd½0 The transformation mix can be prepared in larger volumes and kept at +4°C for up to 24 hours.
13. Resuspend the cells by gently pipetting up and down in 100 μL of 0.2M LiAc + 20% glycerol solution.
14. Mix the 100 μL cell suspension with 100 μL of the transformation mix, 20 μL of digested plasmid, and 240 μL of 58% PEG solution. Pipette up and down or briefly vortex until the contents are well mixed.
15. Incubate at room temperature on a rotating wheel or in a tabletop Thermomixer with gentle shaking (400 rpm) for 30 minutes.
16. Add 54 μL of DMSO to increase DNA uptake efficiency. Mix by briefly vortexing.
17. Incubate at 42°C for 20 minutes in a tabletop Thermomixer with vigorous (950rpm) shaking.
18. Centrifuge for 1 minute at 3000g at room temperature. The cells should form a white deposit on the wall of the tube. Remove the supernatant carefully.
19. Resuspend in 1 mL of YPD and incubate at room temperature for 20 minutes.
20. Centrifuge for 1 minute at 3000g and remove supernatant.
21. Resuspend in 200 μL of TE buffer by pipetting and plate the entire cell suspension onto an SD agar plate lacking the appropriate auxotrophic marker (Synthetic Defined Media Dropout Plate).
22. Incubate the plate upside down at 30°C for 2 days.
23. Streak the original, untransformed strain on a rich media plate (YPD) to get single colonies and incubate upside down at 30°C for 2 days. This will be used as a negative control on Day 3.

### Day 3: Colony PCR

24. Check that single colonies have appeared on the transformation plate. Colonies usually appear within 2 days, but this might take longer depending on the division rate of the strain background and the temperature of incubation.
25. Pick colonies with a clean 10-μL pipette tip and resuspend in a PCR tube containing 4μL of YPD. The transformation efficiency is typically very high (>90%) and, therefore, checking 3-4 colonies by PCR is generally enough to find a correctly transformed colony.
26. Pipette 3 μL of 20 mM NaOH solution into separate PCR tubes, one per colony to test. Then, transfer 2 μL of each colony suspension into the corresponding tube containing the NaOH solution. Store the remaining colony suspensions at +4°C.
27. Resuspend a single colony of the untransformed negative control strain streaked in Step 23 in 3 μL of 20 mM NaOH solution.
28. Heat all of the cell suspensions at 99°C for 10 minutes in a PCR machine. Cool down to 9°C.
29. Prepare the following master mix for as many reactions as the number of colonies selected in Step 25 + two extra reactions. The volumes given are for a single reaction.

a. 5μL 5M Betaine
b. 2.5μL ThermoPol Buffer
c. 2.5μL 2mM dNTP
d. 0.5μL of each 100mM primer (see Table 2 for the appropriate primers to use for each plasmid and its corresponding auxotrophic marker)
e. 1.25 U Taq DNA polymerase
30. Transfer 22μL of the master mix onto each reaction of boiled cells. Vortex briefly.
31. Run the following PCR program on all tubes.

a. 5 min, 94°C.
b. 30s, 94°C.
c. 30s, 58°C.
d. 45s, 72°C.
e. Repeat steps b through d, 30 times in total.
f. 10 min, 72°C.
g. Hold at 9°C.
32. Add loading dye to the completed PCR reactions to a final concentration of 1X and run on a 2% electrophoresis gel in 1X TBE buffer. See Table 2 for the results expected if the plasmid has been correctly integrated.
33. Pick one colony that yielded correct PCR results and start a 5-mL culture in Synthetic Defined (SD) media lacking the auxotrophic marker of the repressor plasmid. Shake at 30°C overnight, 275 rpm.
34. Mix 0.8 mL of the overnight culture with 0.2 mL of 70% glycerol and store at −80°C for future use. Inoculate a small amount from this stock into liquid YPD to revive the strain whenever necessary.

## BASIC PROTOCOL 2

### Generating a WTC_846_ strain with controlled expression of the targeted gene

This multi-day protocol describes how to replace the promoter controlling the expression of a *S. cerevisiae* endogenous, chromosomal gene with the WTC_846_ promoter P_7tet.1_. It can be used for any endogenous gene, except for *TDH3*, for which Alternative Protocol 1 is recommended. Executing this Basic Protocol 2 requires the investigator to decide on which antibiotic resistance marker to use, select the appropriate P_7tet.1_ plasmid for PCR amplification and acquire the plasmid from Addgene ahead of time. We describe how to design primers for generating a PCR product with overhangs for homologous repair using this plasmid. On Day 1, a LiAc-based transformation is carried out as described in Basic Protocol 1 with this PCR product. This transformation requires short strain preparation steps three days (Day −2) and one day (Day 0) before it can be executed. When performed using the parent strain generated in Basic Protocol 1, this resulting WTC_846_ strain allows tightly regulated control of the targeted gene by exogenous anhydrotetracycline or its derivatives.

### Materials

All reagents required for Basic Protocol 1 are also required for Basic Protocol 2. This protocol additionally requires the following reagents:

Hygromycin (CAS number 31282-04-9), dissolved in ddH_2_O

Nourseothricin (CAS number 96736-11-7), dissolved in ddH_2_O

YPD plates with aTc and Nourseothricin or Hygromycin (See Reagents and Solutions)

10 X Taq/Vent Buffer (See Reagents and Solutions)

Vent DNA Polymerase (NEB M0254)

Phusion polymerase (NEB Phusion, M0530, or equivalent high-fidelity polymerase)

Gel purification kit (Quiagen QIAquick Gel Extraction Kit, 28706X4, or equivalent)

Anhydrotetracycline, dissolved in 70% (v/v) Ethanol (CAS number 13803-65-1)

Plasmids from Addgene (See Step 1)

### Protocol steps

1. Decide which antibiotic resistance to use and pick the appropriate tagging plasmid carrying the marker and P_7tet.1_. Two are provided via Addgene (https://www.addgene.org/browse/article/28203500/), P2350 encodes Hygromycin resistance and P2375 encodes Nourseothricin resistance. Order well in advance. *The choice of antibiotic resistance does not make a difference in the functioning of WTC_846_. The strain to be transformed should not already carry a resistance gene for the chosen antibiotic. We normally prefer to use Nourseothricin, if possible, because of its stability. Liquid stocks of concentrated Nourseothricin can be made and kept at −20*°C *for at least a year without any observable loss of activity. Plates can be poured in advance and kept at +4C for up to 12 months. For reasons we have not explored, hygromycin plates stored at +4°C are not stable for more than a month.*
2. Perform steps 2 through 5 described in Basic Protocol 1 to prepare the plasmid. The final plasmid concentration should be at least 100ng/μL.
3. Decide which extended Kozak sequence to use. This is the last 15 bp before the start codon of the controlled gene. It influences translation efficiency, thus affecting the basal uninduced expression of the system. Three Kozak sequences are given below in decreasing order of translation efficiency (sequence in parentheses indicates the reverse complement). It is especially recommended to use the least efficient Kozak sequence for low abundance endogenous proteins. The highest efficiency Kozak is recommended for high abundance proteins and for overexpression.

K1. ACACACATAAACAAA (TTTGTTTATGTGTGT)
K2. AGAGAGAGAGAGAGA (TCTCTCTCTCTCTCT)
K3. AAGGGAAAAGGGAAA (TTTCCCTTTTCCCTT)
4. Design primers to generate a PCR product using the tagging plasmid selected in Step 1 based on the antibiotic resistance to be used. These primers will have portions (3’) that hybridize to the tagging plasmid and tails (5’) to hybridize to the *S. cerevisiae* chromosome. The annealing portions are standardized, whereas the tails will determine which endogenous promoter will be replaced, and how much of the endogenous promoter will be removed in the process. See Figure 4 for a visual representation of the primer designs.

a. Design the forward primer. Use the sequence 5’agcttgccttgtccccgcc3’ as the annealing portion of the primer. Select 40 base pairs upstream of the start codon of the gene to be controlled and use this sequence as the 5’ tail of your forward primer, such that the primer reads: 5’-40bp upstream sequence-annealing portion-3’. *The 40 base pairs chosen within the promoter region of the GOI do not need to be immediately upstream of the start codon. The region between these 40 base pairs and the start codon of the targeted gene will be removed during the transformation-homologous repair process. It is thus possible -though not mandatory- to remove the entire endogenous promoter if the exact sequence is known. However, it is important not to interfere with other upstream endogenous genes, terminators, and promoters in the process, and the choice of which DNA to replace should therefore be made on a case-by-case basis after consideration of the surrounding genome sequence. If it can be done without disturbing upstream genes, we generally recommend removing 100-200bp upstream of the targeted gene to disrupt any naturally occurring regulatory elements that might affect the smooth functioning of the synthetic P_7tet.1_*.
b. Design the reverse primer. Use the sequence 5’tttattcgaaactaagttcttggtg3’ as the annealing portion of the primer. This will anneal to the sequence 5’caccaagaacttagtttcgaataaa3’ on the tagging plasmid. Then, use the reverse complement of the first 40 base pairs of the targeted gene (including start codon), followed by the reverse complement of the chosen Kozak sequence as the 5’ tail. The primer will read as follows: 5’-reverse targeted gene-reverse Kozak sequence-annealing portion-3’.
c. *Alternative reverse primer:* If desired, it is possible to add a flag tag to the open reading frame of the targeted gene using this reverse primer design. (Brizzard & Chubet, 1997; Hopp et al., 1988). These tagging plasmids provide the option of generating a fusion protein containing an N-terminal flag tag followed by an 8-glycine linker to the protein encoded by the targeted gene. In this case, the Kozak sequence cannot be altered and the high efficiency Kozak sequence present on the tagging plasmid will be used. For this option, use the sequence 5’catcgatgaattctctgtcgg3’ as the annealing portion of the primer. This will anneal to the S4 primer binding site on the tagging plasmid (5’ccgacagagaattcatcgatg3’). Use the reverse complement of the first 40 base pairs of the controlled gene (excluding the start codon) as your 5’ tail. The primer will read as follows: 5’-reverse controlled gene sequence-annealing portion-3’.
5. Perform a PCR using the P_7tet.1_ promoter-replacing (“tagging”) plasmid chosen in Step 1 as the template and the forward and reverse oligos designed in Step 3. Use any high-fidelity polymerase protocol and appropriate annealing temperatures (T_a_), or the PCR protocol below (adapted from (Janke et al., 2004)) and a T_a_ of 48°C (for the reverse primer designed in 3b) or 50°C (for the reverse primer designed in 3c).

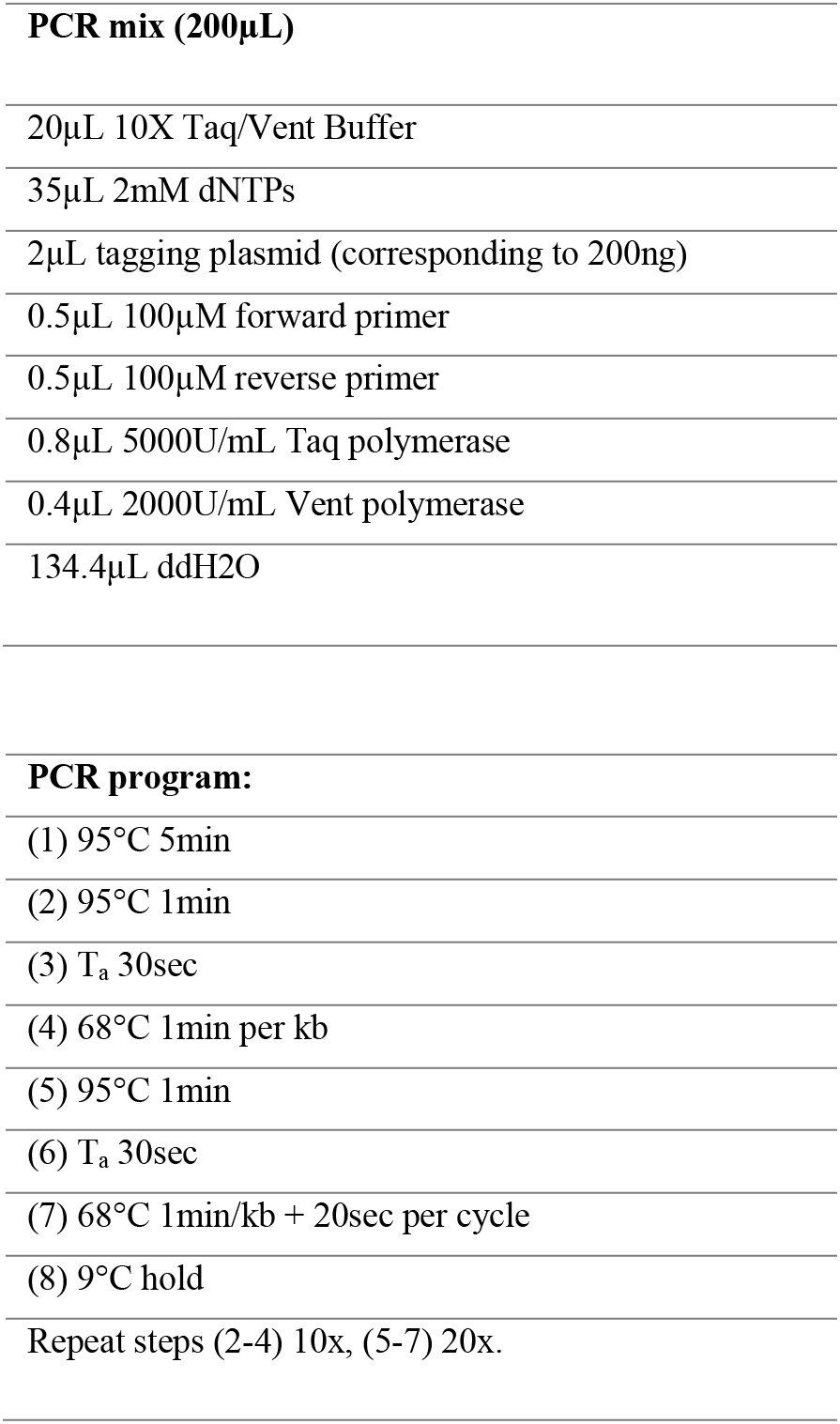 *The resulting PCR product will be 2463bp long with the reverse primer designed in Step 3b, or 2597bp long with the reverse primer designed in Step 3c.*
6. Run the PCR product on a 1% electrophoresis gel. Isolate and gel purify the P_7tet.1_ promoter replacing tagging PCR product and resuspend in ddH_2_O at a concentration of around 50ng/μL.
7. Design colony PCR primers for checking correct integration of P_7tet.1_ in front of the targeted gene.

a. Use the sequence 5’cagttcgagtttatcattatcaatactg3’ as the forward primer. This anneals to the start of P_7tet.1_.
b. Design a reverse primer with a compatible annealing temperature (T_a_) that anneals close to the start of the targeted gene. The 5’ and 3’ ends can anneal anywhere inside the Open Reading Frame of the targeted gene, as long as the expected PCR product is less than 1kb. *Note that this colony PCR strategy can be used to check for correct replacement of any promoter of any endogenous gene (Gene Of Interest, or GOI), except for P_TDH3_ of S. cerevisiae, from which P_7tet.1_ was derived. Since P_7tet.1_ is highly homologous to P_TDH3_, the primers designed here will always give a positive PCR result, regardless of correct promoter replacement. If P_TDH3_ replacement is desired, always use Alternate Protocol 1 and design a separate forward primer that would hybridize to the TetR binding sites within P_7tet.1_, and therefore would not hybridize to P_TDH3_*

### Day −2 to Day 0: Strain Preparation for Yeast Transformation

8. **Day −2:** Streak the parent yeast strain generated in Basic Protocol 1 onto a YPD plate and incubate at 30°C for 2 days.
9. **Day 0:** Start a liquid culture of the parent yeast strain to be transformed by inoculating a single colony in 5 mL of rich media (YPD) in a glass culture tube. Shake overnight at 30°C, 275 rpm.

### Day 1: Transformation: Creating the WTC_846_::GOI strain

10. Follow steps 9 through 23 of Basic Protocol 1 with the following changes:

a. In Step 8, use 20μL of the purified P_7tet.1_ tagging PCR fragment solution instead of the plasmid restriction digestion.
b. In Step 19, resuspend in 1 mL of YPD + an appropriate concentration of aTc depending on the targeted gene. See Strategic Planning for how to estimate an appropriate aTc concentration for different genes. Incubate at room temperature (~20-22°C) on a rotating wheel or in a tabletop Thermomixer with gentle oscillation (400 rpm) for 3 to 4 hours.
c. In Step 21, plate the entire suspension on YPD + Nourseothricin or Hygromycin + aTc, as appropriate. See Strategic Planning to estimate the appropriate plate concentration of aTc to be used. The concentration needed for a given protein dosage will generally be higher in solid medium than in liquid medium, likely due to slow diffusion of aTc into the growing colony and depletion of aTc from the solid medium immediately adjacent to the colony.

### Day 3: Colony PCR

11. Check that single colonies have grown on the transformation plate. *Colonies usually appear within 2 days, but might take longer depending on the division rate of the strain background and the temperature of incubation. If the endogenous gene is expressed in a highly cell cycle specific manner (e.g. histones), or if the the aTc concentration in the plate results in expression of the controlled gene that is higher or lower than the endogenous expression level, this difference in protein dosage may also slow down appearance of colonies. See Strategic Planning and Troubleshooting*.
12. Pick colonies with a clean 10-μL pipette tip and resuspend in a PCR tube containing 4μL of YPD + an appropriate amount of aTc (determined as explained in Strategic Planning). *The transformation efficiency is low, especially with essential genes (~10%) and therefore checking at least 10 colonies using PCR is recommended. See Alternative Protocol 1 for achieving higher efficiency.*
13. Pipette 3 μL of 20 mM NaOH solution into separate PCR tubes, one per colony to test. Then, transfer 2 μL of each colony suspension into the corresponding tube containing the NaOH solution. Store the remaining colony suspensions at +4°C.
14. Resuspend a single colony of the untransformed negative control strain in 3 μL of 20 mM NaOH solution.
15. Heat all of the cell suspensions at 99°C for 10 minutes in a PCR machine. Cool down to 9°C.
16. Prepare the following master mix for as many reactions as the number of colonies selected in Step 12 + two extra reactions. The volumes given are for a single reaction.

a. 5μL 5M Betaine
b. 2.5μL ThermoPol Buffer
c. 2.5μL 2mM dNTP
d. 0.5μL of each 100mM primer (use primers designed in Step 7)
e. 1.25 U Taq DNA polymerase
17. Transfer 22μL of the master mix onto the boiled cells. Vortex briefly.
18.Run the following PCR program on all tubes.

a. 5 min, 94°C.
b. 30s, 94°C.
c. 30s, Ta°C.
d. 45s, 72°C.
e. Repeat steps b through d, 30 times in total.
f. 10 min, 72°C.
g. Hold at 9°C.
19. Run the completed PCR on a 1% electrophoresis gel. The PCR product spans the entire P_7tet.1_ promoter and includes a portion of the open reading frame of the GOI. The length of this portion depends on where the reverse primer designed in Step 7 anneals. Therefore, the expected length of the PCR product will be the total length of P_7tet.1_ (730bp) and the length between the start codon and the 5’ end of the reverse primer. *Even though incorrect integration is extremely rare, it is still advisable to gel isolate and submit this PCR product for Sanger sequencing at a sequencing facility and to subsequently inspect the resulting sequence to verify that the entire P_7tet.1_ sequence has been correctly integrated.*
20. Pick one colony that yielded correct PCR results and start a 5-mL culture in YPD+appropriate antibiotic+appropriate aTc concentration. Shake at 30°C overnight. *If the PCR product from step 19 was sent to sequencing, the user can pick more than one colony and stock these in the next step, to keep until the results of the Sanger sequencing are available.*
21. Mix 0.8mL of the overnight culture with 0.2mL 70% glycerol and store at −80°C for future use. Inoculate a small amount from this stock into liquid YPD+appropriate aTc concentration with or without the antibiotic to revive the strain whenever necessary. *The promoter is now stably integrated and it is not mandatory to keep this strain under antibiotic selection to maintain WTC_846_ functionality.*

## ALTERNATE PROTOCOL 1

### CRISPR-mediated promoter replacement

This multi-day protocol describes the steps to replace the promoter of any targeted *S. cerevisiae* gene with the WTC_846_ promoter P_7tet.1_ using a CRISPR-Cas9-mediated double strand break (Figure 3), with or without an antibiotic marker. While it takes 3 days longer than Basic Protocol 2, making this break enhances the efficiency of correct homology repair-directed integrations (Figure 7) and the number of recovered correctly integrated transformants. With this approach, the investigator can also decide to generate “markerless” strains that do not include an antibiotic-resistance marker. The protocol involves designing single guide oligos for targeting the Cas9 enzyme to the desired cut site and carrying out a transformation to assemble the CRISPR-Cas9 expression plasmid inside the yeast cell. This first step creates a strain we call the CRISPR Mid Strain, which is capable of cutting the genomic DNA at the targeted site but does not have the repair template. A second transformation introduces a PCR product as a repair template such that the double-stranded cut generated by Cas9 can be repaired through homology repair.

**Figure 7.**
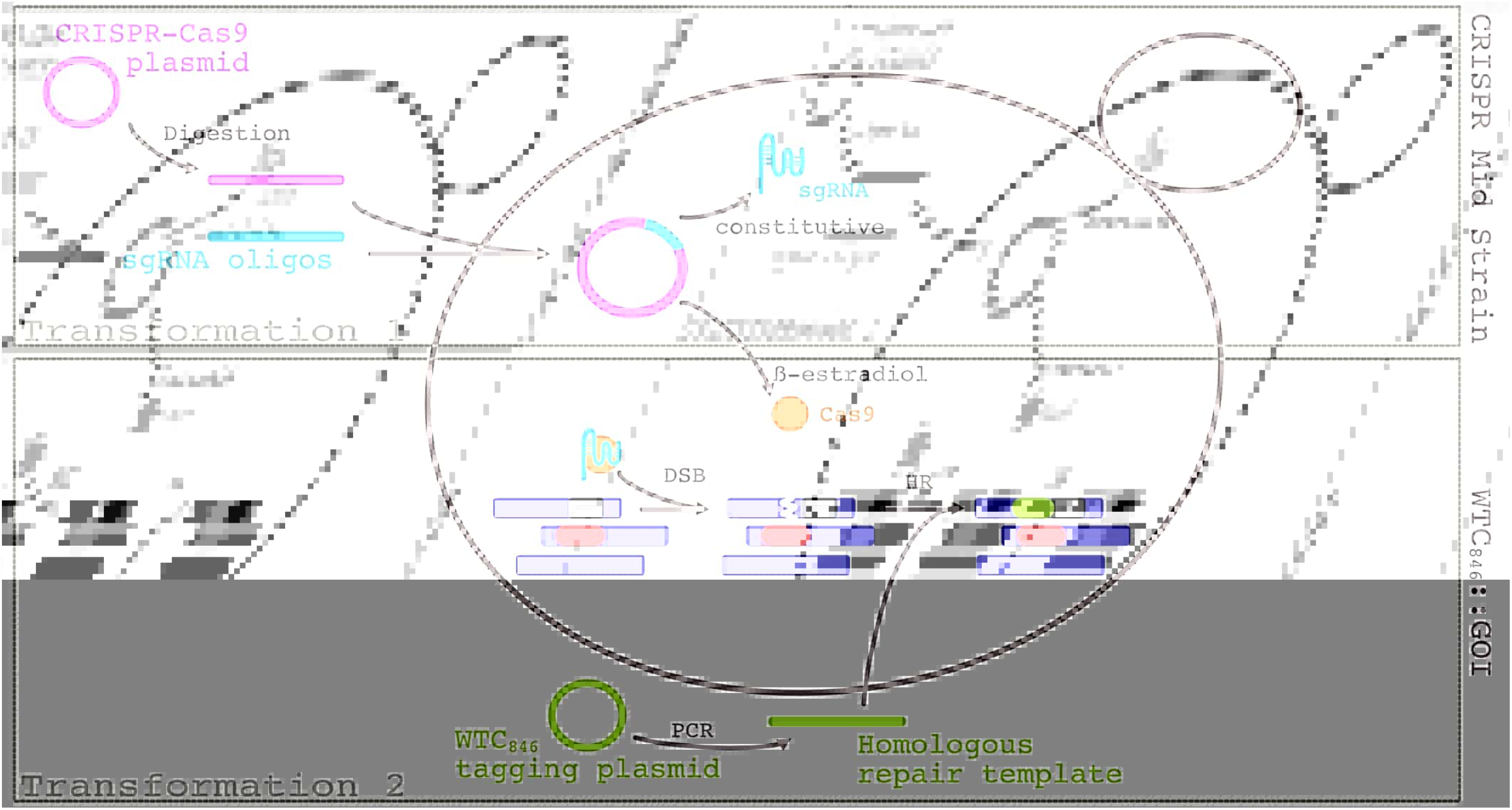
Overview of the CRISPR-Cas9 based transformation protocol (Alternate Protocol 1). The two transformation steps required to replace the promoter of an endogenous gene are depicted. Transformation 1 involves linearizing the CRISPR-Cas9 plasmid and co-transforming it together with a double stranded sgRNA oligo. These assemble inside the cell to generate the CRISPR Mid Strain. This strain is capable of generating a double stranded break at the target genomic locus upon induction of Cas9 expression by β-estradiol but cannot repair it, as it does not possess the correct repair template. The second transformation is performed on this strain to introduce the repair template. Cas9 expression is induced using 10μM β-estradiol. Cas9 then complexes with the constitutively expressed sgRNA to target a region within the GOI promoter. This creates a double stranded break in the DNA. The cell is simultaneously transformed with a PCR product that contains the P_7tet.1_ promoter and homology arms to the region where the double stranded break was generated. The result is a strain where WTC_846_ controls the expression of the GOI.

Additionally, this protocol is necessary for the construction of strains in which two or more genes are to be brought under the control of WTC_846_. Such constructions require serial transformations to replace different promoters with P_7tet.1_. Unless the investigator makes a targeted break for the genes to be controlled after the first promoter replacement, it is more likely that the repair template will recombine with the already existing P_7tet.1_ rather than replacing the promoter of the second targeted gene. The double stranded break induced by Cas9 in this protocol increases the likelihood that a correct replacement is achieved.

### Additional Materials to Basic Protocol 2

All reagents required for Basic Protocols 1&2 are required for Alternate Protocol 1. This protocol additionally requires the following reagents:

BsaI restriction enzyme (NEB, R3733)

AflII restriction enzyme (NEB, R0520)

YPD plates with G418 (See Reagents and Solutions)

YPD media with G418 (See Reagents and Solutions) β-estradiol (CAS number 50-28-2), dissolved in ddH_2_0 Plasmid P2061 from Addgene (contains β-estradiol inducible Cas9, https://www.addgene.org/186299/)

### Protocol steps

1. Follow Basic Protocol 2, steps 1-6 if promoter replacement will be done using an integrated antibiotic marker.
2. If a markerless promoter replacement is desired, follow Basic Protocol 2, Steps 1-6 with the exception of 4a. Instead design the forward oligo as follows:

a. Use the sequence (5’cagttcgagtttatcattatcaatactg3’) as the annealing portion of the primer. Select 40 base pairs upstream of the start codon of the GOI and use this sequence as the 5’ tail of your forward primer such that the primer reads: 5’-40bp upstream sequence-annealing portion-3’. The region between these 40 base pairs and the start codon of the controlled gene will be removed during the transformation process. It is advisable to remove as much of the endogenous promoter as possible when not using an antibiotic marker. The goal in removing the endogenous promoter is to eliminate any endogenous regulation or activity, such as silencing mediated by the endogenous promoter, or transcription that might originate from the endogenous promoter upstream of P_7tet.1_, and cause transcriptional interference. However, it is important not to interfere with other endogenous genes, terminators and promoters in the process.
3. Design colony PCR oligos for checking correct integration by following Basic Protocol Step 7.
4. Design single guide RNA (sgRNA) oligos for generating a cut site within the promoter to be replaced.

a. Choose a position upstream of and in proximity to the gene, where any base (N) is followed by two guanine bases (NGG site). This NGG site needs to be mutated at least at one or both G residues, or removed by the desired promoter replacement, such that only in correctly transformed cells would Cas9 be unable to induce further double-stranded breaks. Therefore, pick this NGG site to be within the region that will be removed once the homology-directed repair template generated through PCR in Steps 2 or 3 are integrated into the genome.
b. Design the sgRNA sequence as 5’gcgccggctgggcaacaccttcgggtggcgaatgggacc[20nt]gttttagagctagaaatagcaagttaaaataaggc3’, using the 20 nucleotides immediately upstream of the chosen NGG.
c. Order forward and reverse (i.e., complementary) oligos of the final sgRNA at genomics scale.
d. To anneal the two complementary sgRNAs, resuspend them at 100mM concentration, mix 7 μL of each in a 0.2-mL tube, heat to 95°C for 5 minutes, and let cool to room temperature.

### Day −2 to Day 0: Preparation for Yeast Transformation

5. **Day −2:**Streak the parent yeast strain generated in Basic Protocol 1 onto a YPD plate and incubate at 30°C for 2 days.
6. **Day 0:** Start a liquid culture of the parent yeast strain to be transformed by inoculating a single colony in 5 mL of rich media (YPD) in a glass culture tube. Shake overnight at 30°C, 275 rpm.
7. **Day 0:** Linearize 1 μg P2061 in a restriction enzyme digestion using the enzymes BsaI and AflII simultaneously. Follow the enzyme manufacturer’s instructions. Run on a 0.5% electrophoresis gel and perform a gel clean up (Voytas, 2001). The gel run and cleanup are not mandatory but recommended at least once to make sure the enzymes are linearizing the plasmid.

### Day 1. First Transformation: Creating the CRISPR Mid Strain

The CRISPR Mid Strain is a strain where the plasmid that will express Cas9 and the targeting sgRNAs have been assembled (Figure 7). This strain will create a double stranded break at the targeted genomic locus upon induction with β-estradiol, but cannot repair it correctly since it does not have a repair template available. This repair template is introduced via a second transformation on Day 5.

8. Perform two transformations: one assembly transformation with the digested CRISPR-Cas9 plasmid (P2061) and the double stranded sgRNA, and one negative control transformation with only the digested P2061. Follow Basic Protocol steps 9 through 22 with the following modifications.

a. In Step 14, add 50 ng of linearized P2061 to both transformations, and 10 μL of the sgRNA mixture (designed in this protocol, Step 4b) to the assembly transformation. Omit the sgRNA mixture from the negative control transformation.
b. In Step 19 incubate at room temperature for 3 to 4 hours.
c. In Step 21 plate on YPD+G418 plate.

### Days 3 & 4: Spotting Assay

9. Check that single colonies have appeared on the plates. It is possible that no colonies will appear on the negative control plate., As long as the assembly transformation has many more colonies than the negative control plate, assume the assembly has worked correctly.
10. Resuspend 5 colonies from the assembly transformation and 1 colony from the control transformation in 100μL YPD+G418. Create six 1:10 serial dilutions in the same media and spot 4μL of each dilution onto two YPD plates, one containing G418 and the other G418 + 10μM β-estradiol. Incubate at 30°C overnight. β-estradiol induces Cas9 expression. In colonies that have correctly assembled the CRISPR-Cas9 plasmid together with the provided sgRNA, this will lead to simultaneous Cas9 and sgRNA expression, which will induce a double-stranded DNA break. Without a repair template, continuous DNA damage will slow down growth. Therefore, there will be a visible difference between the growth rate of correctly transformed colonies between the G418 and G418+ β-estradiol plates.
11. Compare the two plates with and without β-estradiol to see which colonies grow slower in the presence of β-estradiol. See Figure 8 for an example.
12. Pick one correctly transformed colony from the plate with G418 and without β-estradiol and use it to start an overnight culture in YPD+G418. It is important to start the culture from the plate without β-estradiol to avoid potential mutations introduced by the continuous DNA damage caused by Cas9. This strain should always be grown in the presence of G418; otherwise, it will lose the assembled CRISPR-Cas9 plasmid quickly.

**Figure 8.**
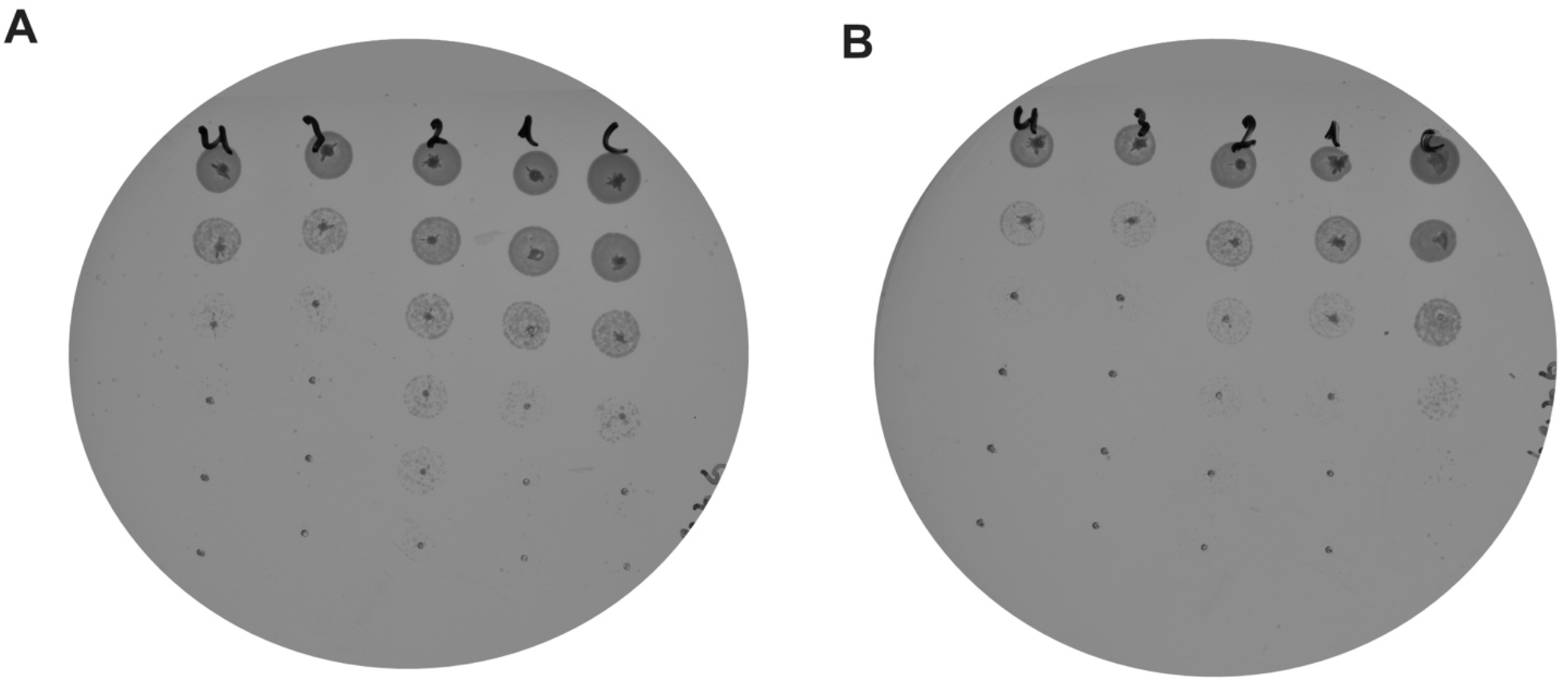
An example of a successful spotting assay after CRISPR-Cas9 plasmid assembly. A CRISPR Mid strain was generated as described in Alternate Protocol 1 and depicted in Figure 7. Four single colonies were selected from the selection plate of the assembly transformation (with sgRNA oligos, numbered 1-4), and one single colony was selected as control from the negative control transformation (without sgRNA oligos, labelled c). These were spotted onto YPD+G418 plates without (A) and with (B) 10μM β-estradiol. The top row has the highest concentration of cells and each row is a 1:10 dilution. Growth rate differences between the two plates are evident when the number of colonies in each row is compared. All colonies 1-4 show slower growth on the β-estradiol plate, as opposed to the control colony which shows equal growth on both plates.

### Day 5: Second Transformation

13. Prepare a glycerol stock of the CRISPR Mid Strain by mixing 0.8 mL of the overnight culture with 0.2 mL of 70% glycerol. Store at −80°C for future use. Always revive this strain in presence of G418.
14. Dilute the overnight culture into 10 mL of YPD+G418 in a glass culture tube. If no cell counter is available, dilute down to an OD_600_ of 0.5 or just use a 1:50 dilution ratio (i.e. add 200 μL to the 10-mL culture). If one is available, dilute the culture to a concentration of 2.5 million cells/mL. Shake at 30°C for 3 hours at 275rpm until the culture reaches a concentration of 10 million cells/mL or an OD_600_ of 2.0.
15. Add β-estradiol to the culture at a final concentration of 10μM about 1.5 hours before the desired cell density is reached. Assuming doubling every 1.5h hours, this will be when the culture reaches a density of 5 million cells/mL or an OD_600_ of 1.0.
16. For a replacement using an antibiotic marker, perform the transformation by following Basic Protocol 1, Steps 9-23 with the following modifications.

a. In Step 14, add 500 ng of the PCR template generated in Step 1 of this protocol, instead of a plasmid digestion.
b. In Step 19, incubate at room temperature for 3-4 hours in YPD+ G418+ 10μM β-estradiol+ an appropriate amount of aTc for the GOI. See Strategic Planning for how to estimate appropriate aTc concentrations for different genes.
c. In Step 21 plate the entire suspension on YPD + G418 + the new antibiotic+ aTc. See Strategic Planning to decide on the aTc concentration necessary.
17. For a markerless replacement, perform the transformation by following Basic Protocol 1 Steps 9-23 with the following modifications.

a. In Step 14 add 500 ng of the PCR template generated in this protocol, Step 2 instead of a plasmid digestion.
b. In Step 19, incubate at room temperature for 3-4 hours in YPD+G418+ 10μM β-estradiol+ an appropriate amount of aTc for the GOI. See Strategic Planning for how to estimate appropriate aTc concentrations for different genes.
c. In Step 21, plate 1/4^th^ of the cell suspension on YPD + G418 + 10μM β-estradiol + aTc. See Strategic Planning to decide on the aTc concentration necessary.

### Day 7&8: Colony PCR and stocking

18. Follow Basic Protocol 2, Steps 12-19 for finding a correctly transformed colony via PCR using the primers designed in this protocol, Step 3. The efficiency will be higher compared to Basic Protocol 2, so fewer colonies may be screened. If the transformation plate is too crowded to pick single colonies, restreak onto a new plate and incubate for 1-2 days at 30°C until single colonies appear.
19. Once a correct colony is found, grow in liquid YPD with the appropriate amount of aTc (+ the appropriate antibiotic if a marker was inserted) but without G418 overnight, to lose the CRISPR-Cas9 plasmid. It is best to check for complete loss of the CRISPR-Cas9 plasmid if further transformations with G418 resistance as the selectable marker are planned. To do this, streak the overnight culture on YPD+aTc plates for single colonies. Select 4-5 single colonies and streak on YPD+G418+aTc and YPD+aTc plates to ensure that they have lost the plasmid (no growth should be observed on the plate with G418).
20. Mix 0.8mL of the overnight culture with 0.2mL 70% glycerol to create a glycerol stock and store at −80°C. Revive whenever necessary by inoculating into YPD + the appropriate amount of aTc (+ antibiotic marker if used). The promoter is now stably integrated and it is not mandatory to keep this strain under antibiotic selection to maintain WTC_846_ functionality.

## BASIC PROTOCOL 3

### Cell cycle synchronization/Arrest and Release using the *WTC_846-K3_::CDC20* strain

This protocol is intended to demonstrate use of WTC_846_ to induce and turn off expression of a targeted gene. As an example, we use this system to arrest cell cycle progression and then to synchronously release those cells from arrest. In *S. cerevisiae*, such arrest & release experiments are commonly used when investigating proteins hypothesized to function only during certain cell cycle phases(Angeles Juanes, 2017). A synchronized population allows reliable batch measurements of transcript or protein levels, phosphorylation states and similar readouts that might reveal protein function or regulation(Cosma et al., 1999; Ewald et al., 2016). However, there are currently no available methods to completely synchronize batch cultures under different growth conditions. This protocol is described for a Nourseothricin resistant *WTC_846-K3_::CDC20* strain created using a BY4742 background, following Basic Protocols 1 and 2, which can be synchronized in batch culture independent of media conditions. The oligos used to construct this strain are provided in Table 4. *CDC20* is an essential gene regulating the transition from metaphase to anaphase (Pesin & Orr-Weaver, 2008). Upon depletion of *CDC20*, cells arrest in G2/M phase of the cell cycle, and exhibit large buds and 2n DNA content. This strain carries the least efficient extended Kozak given in Basic Protocol 2 (AAGGGAAAAGGGAAA), and thus presumably the lowest protein expression, which is needed for the null phenotype that results in complete arrest. While this protocol describes the assay in complete medium YPD, it can also be performed in any other media.

**Table 4.**
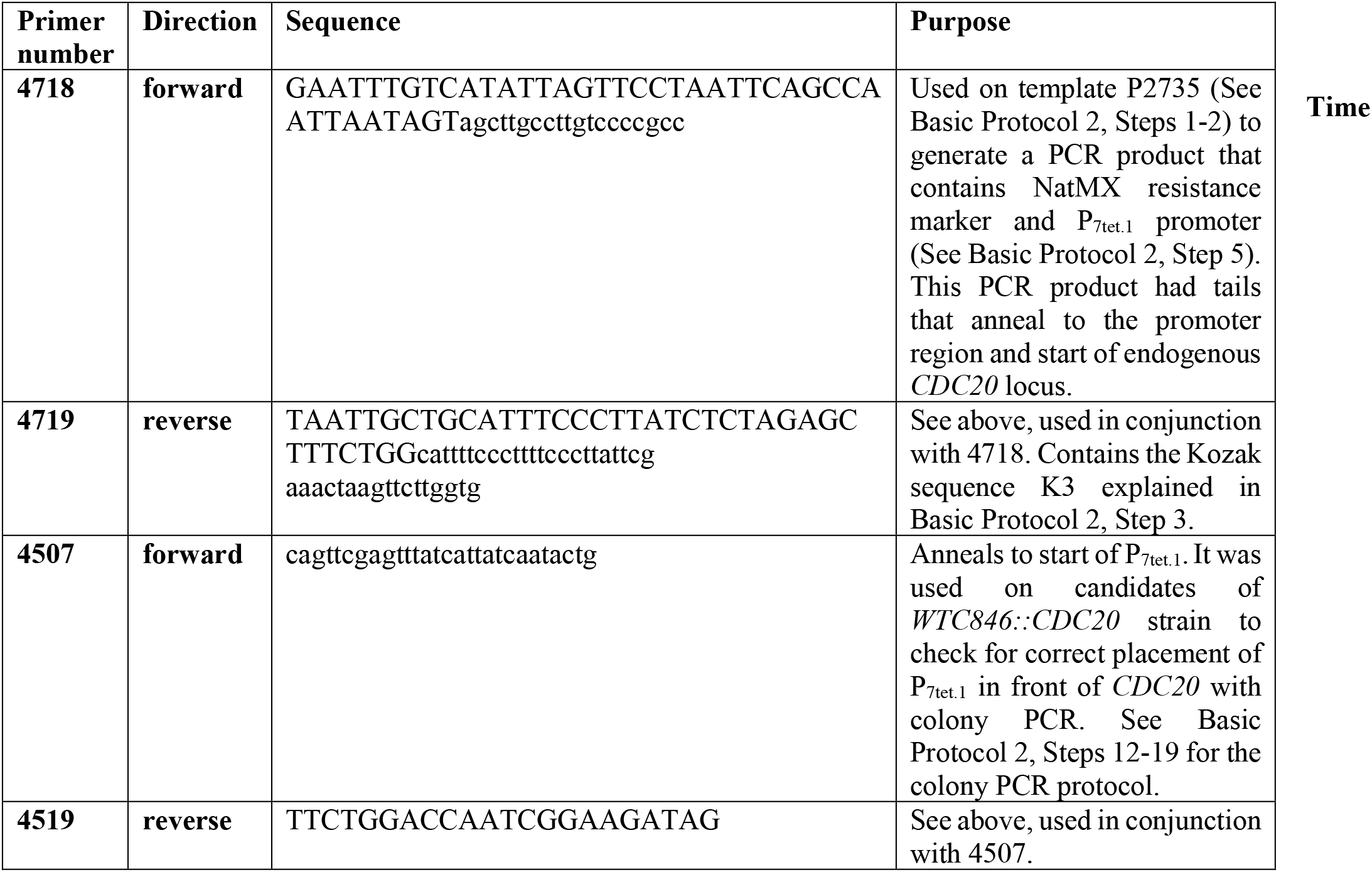
Primers used to generate a *WTC846::CDC20* strain. Primers used to generate the PCR product for homologous repair, and primers used for colony PCR are given for the strain used in Basic Protocol 3.

### Materials

YPD media (See Reagents and Solutions)

YPD media with Nourseothricin (See Reagents and Solutions)

Anhydrotetracycline, dissolved in 70% (v/v) Ethanol (CAS number 13803-65-1)

WTC_846-K3_::*CDC20* strain (can be constructed using the protocols described above or requested from the authors) Shaking incubator (Kuhner Shaker ISF1-X or equivalent)

Cell counter (Beckman, Z2 Coulter or equivalent)

Eppendorf Tubes (1.5 or 2mL)

Glass culture tubes (55mL, Huberlab 9.6131.33 or equivalent)

Tabletop centrifuge (Eppendorf 5514R or equivalent)

Light microscope (Nikon Eclipse E100 or equivalent)

### Protocol steps

#### Day 0

1. Start a culture of the WTC_846_-K3*::CDC20* strain in 4mL YPD + Nourseothricin + 5ng/mL aTc. Shake at 30°C overnight, 275rpm. The inclusion of Nourseothricin is not mandatory but is useful for avoiding contamination of the medium with bacterial or wild type yeast. The assay can be performed in media other than YPD, or at temperatures other than 30°C, however, this will likely change the doubling time of the yeast and, therefore, the growth rate. Bear in mind that the slower the growth rate, the longer it will take all the cells in the population to deplete their pool of Cdc20 by dilution and degradation, reach metaphase, and arrest.

#### Day 1

2. Dilute the overnight culture into 5mL YPD + Nourseothricin + 3ng/mL aTc. If a cell counter is available, dilute to a concentration of 2.5 million cells/mL, otherwise dilute to an OD_600_ of 0.5 or use a 1:50 dilution. Shake at 30°C, 275rpm for 4 hours or until the culture reaches the desired density of 10-20 million cells/mL or an OD_600_ of about 2.0.
3. Once the desired culture density is reached, count cell numbers. Transfer 2.5 million cells into an Eppendorf tube. If no cell counter is available, dilute the culture to an OD_600_ of 0.5 and transfer 1 mL to an Eppendorf tube. Spin down at 1000g for 1 minute at room temperature.
4. Carefully remove all of the supernatant and resuspend in 1mL YPD. Repeat twice but resuspend in 100μL YPD during the last repetition. The wash steps should be performed carefully, leaving minimal residual media behind. It is important to wash out all of the anhydrotetracycline in order to reach a complete arrest.
5. Transfer the cells into a 55 mL glass culture tube containing 5mL YPD + Nourseothricin and shake at 30°C for 3-4 hours, 275rpm. Resuspension in YPD without aTc starts the arrest phase of the experiment. No further CDC20 will be produced. Cells will continue to grow for a few cell divisions and this will dilute the existing CDC20 pool. Arrest will be seen once the existing CDC20 protein product is depleted.
6. Confirm arrest using light microscopy. All cells in the field of vision at 40X magnification should have large buds. See Figure 9 for an example. Examine multiple fields of view and if unbudded cells or cells with small buds are seen, continue shaking and check every 30 minutes. Complete arrest should be reached latest within 5 hours, but will depend on how well the wash steps were performed (any residual aTc will allow cells to continue producing CDC20 at low levels), the doubling time of the yeast cells in the particular media used (the slower the growth, the longer it will take to dilute the existing supply of CDC20), and the initial aTc concentration the sample was cultured in at Step 1 (the higher the initial aTc concentration, the more molecules of Cdc20 in the pool to be diluted).
7. If the culture becomes denser than 30 million cells/mL (OD600 of about 6.0), dilute 1:6 in YPD + Nourseothricin and continue checking for arrest under the microscope every 30 minutes. The appropriate culture density will depend on the specifics of the experiment. For example, if the downstream readout method is protein quantification with western blotting, a high number of cells might be required. In such a case, the culture should always be diluted minimally until arrest is reached. However, in all cases, the culture should always be kept in exponential growth phase until arrest is complete, to ensure that the existing CDC20 pool is being diluted efficiently.
8. Once arrest is complete, add aTc to the culture at a final concentration of 600ng/mL and continue shaking at 30°C, 275rpm. The entire culture will synchronously enter G1 phase within 30-35 minutes. See Figure 10 for an example arrest and release assay where cell cycle phase was determined based on DNA content(Azizoğlu et al., 2021). *This high concentration of aTc is used to ensure release from arrest occurs quickly and uniformly. It would lead to overexpression of CDC20 once the expression level reaches steady state, If necessary, lower levels of aTc can also be used to perform the release and achieve endogenous expression levels of CDC20.*

**Figure 9.**
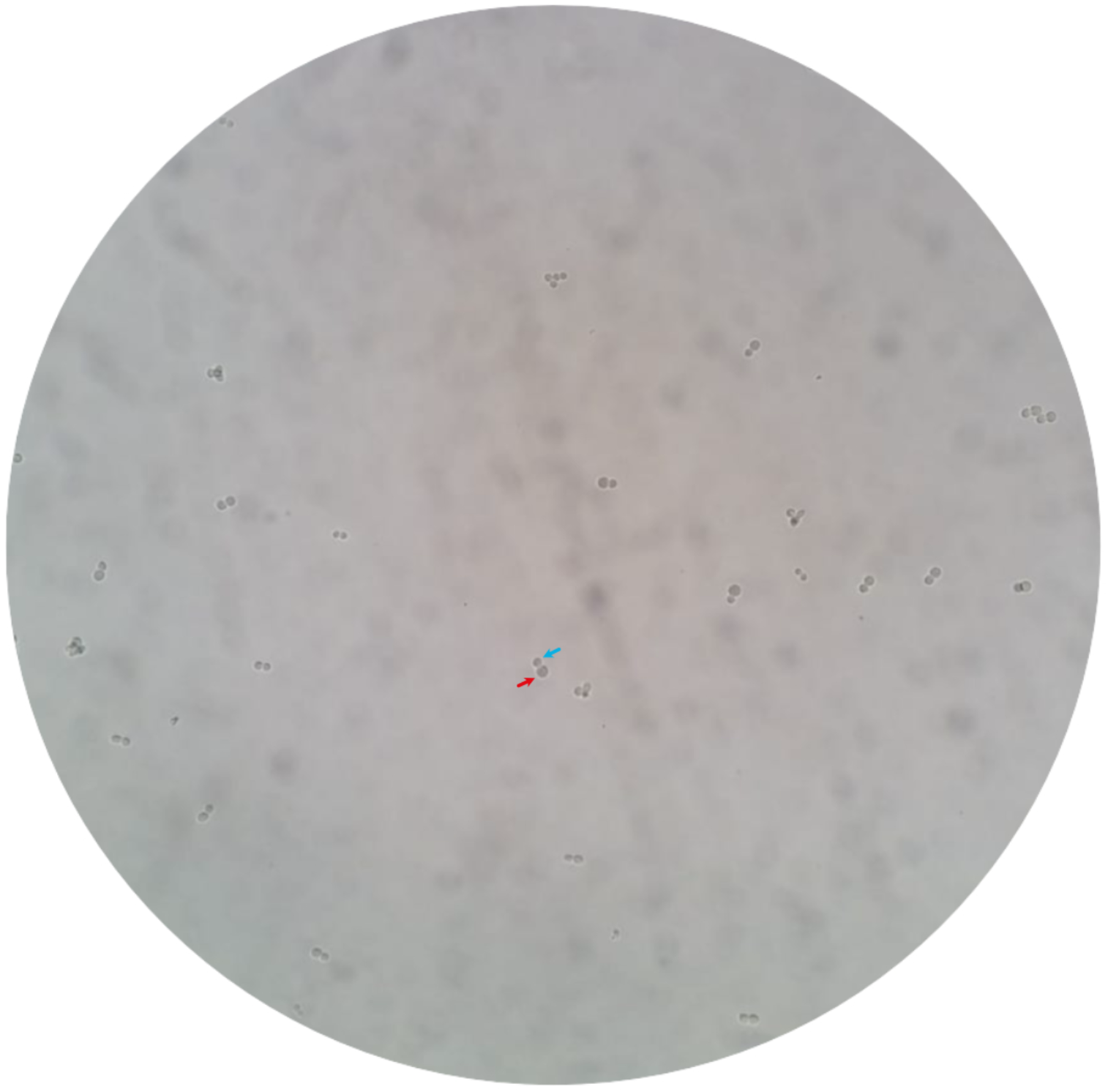
G2/M arrested *WTC_846-K3_::CDC20* cells. Cells were grown in YPD in the presence of 5ng/mL anhydrotetracycline (aTc) overnight such that they would be in exponential phase in the morning. 2.5 million cells were collected, washed, and resuspended in YPD without inducer. 7 hours later, the image above was acquired using a cell phone camera and a light microscope at 40X magnification. The red arrow indicates a mother cell and the blue arrow indicates the large bud (soon to be daughter cell) the mother has produced, but is unable to detach from due to the cell cycle block. The uniform presence of large buds is indicative of G2/M arrest.

**Figure 10.**
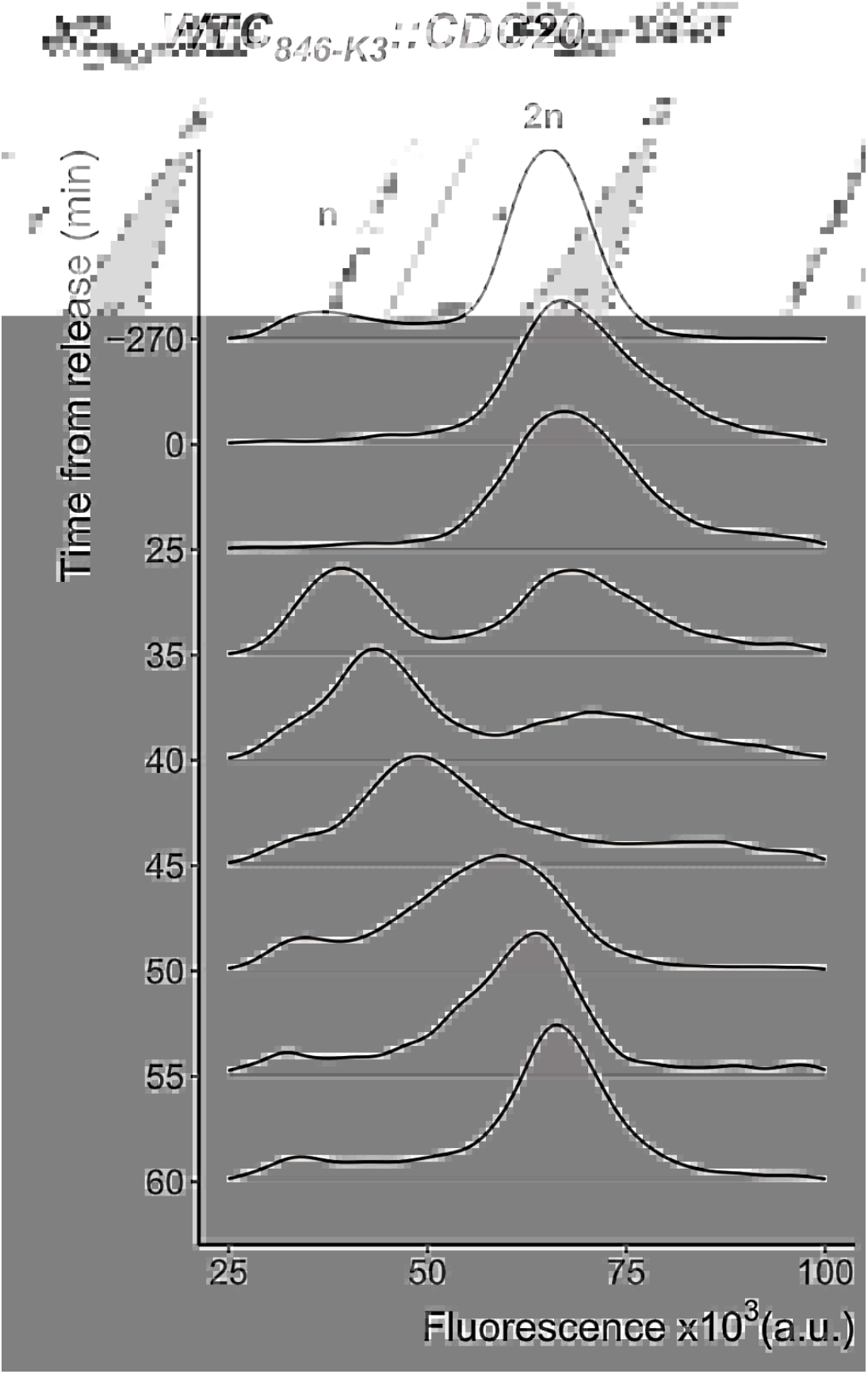
Synchronization of *WTC_846-K3_::CDC20* cells. The figure shows data from flow cytometry. A batch liquid culture of WTC_846-_*_K3_::CDC20* cells growing in 20ng/mL aTc was arrested by aTc withdrawal. After complete arrest, cells were released by addition of 600ng/mL aTc at time 0. Samples were taken at indicated times, fixed in ethanol, and stained with Sytox Green for DNA content analysis using flow cytometry (488nm excitation, 515/20nm emission). The plots are density distributions of a population of 10000 cells per time point, such that the area under the curve equals 1. The fluorescence peaks corresponding to one and two sets of chromosomes, indicating G1 and G2/M phases, are labelled. Figure was reproduced with permission from (Azizoğlu et al., 2021).

## REAGENTS AND SOLUTIONS

### YPD media/plates

- For YPD media, dissolve 1% (w/v) Yeast Extract (Gibco Bacto Yeast Extract or equivalent, CAS no. 8013-01-2), 2% (w/v) Peptone (Gibco Bacto Peptone or equivalent, CAS no. 73049-73-7), and 0.003% (w/v) Adenine (CAS number: 321-30-2) in double distilled water. For YPD plates, also include 2% (w/v) Agar (Cas Number 9002-18-0).
- In a separate container, prepare a 20% (w/v) glucose (CAS number: 50-99-7) solution in double distilled water.
- Autoclave both solutions.
- Add 20% Glucose to the first solution at a final concentration of 2% (v/v).
- If needed, add antibiotics and anhydrotetracycline to both liquid and solid media after autoclave and only once the solution has cooled down to at least 60°C. Use Hygromycin (CAS number 31282-04-9) at a final concentration of 200 μg/L, Nourseothricin (CAS number 96736-11-7) at a final concentration of 100 μg/L, and Geneticin (G418 Sulfate, CAS number 108321-42-2) at a final concentration of 350μg/mL.
- For liquid media without anhydrotetracycline, store at room temperature for up to 3 months. If anhydrotetracycline was added, store at +4°C for up to a month and keep away from light.
- Store agar plates without anhydrotetracycline upside down and sealed in a plastic bag at +4°C for up to 12 months. If anhydrotetracycline was added, store at +4°C for up to a month and keep away from light.

### LB Plates with Carbenicillin

- For LB media, dissolve 0.5% (w/v) Yeast Extract (Gibco Bacto Yeast Extract or equivalent, CAS no. 8013-01-2), 1% (w/v) Tryptone (Gibco Bacto Tryptone or equivalent, CAS no. 91079-40-2), 1% (w/v) NaCl (Merck NaCl or equivalent, CAS no. 7647-14-5) in double distilled water. For LB plates, also add 1.5% (w/v) Agar (CAS no. 9002-18-0).
- Autoclave the solution.
- Add Carbenicillin (Biochemica Carbenicillin Disodium Salt or equivalent, CAS no. 4800-94-6) to a final concentration of 100μg/ml after autoclave and after the solution has cooled down to at least 60°C.
- Store agar plates upside down and sealed in a plastic bag at +4°C for up to 12 months.

### Tris-HCl Buffer

- Dissolve Tris (CAS number: 77-86-1) to final concentration of 0.4M in double distilled water.
- Adjust pH to 6.8 using HCl (CAS number: 7647-01-0) in a fume hood.
- Filtrate with a tabletop filtration system (Merck Steritop or equivalent with 0.22μm pore size) to sterilize.
- Store at room temperature for up to 24 months.

### 0.08M EDTA solution

- Dissolve EDTA (CAS number: 6381-92-6) to final concentration of 0.08M in double distilled water.
- Add NaOH pellets (CAS number: 1310-73-2) to adjust pH to 8.
- Sterilize by autoclavation.
- Store at room temperature for up to 24 months.

### Salmon Sperm DNA solution

- Dissolve ssDNA (CAS number 9007-49-2) in TE buffer (0.1M Tris-HCl buffer, 0.01M EDTA solution) to a final concentration of 1%(w/v) in a 50 mL Falcon tube.
- Dissolve by gentle shaking or inversion at +4°C overnight.
- To sterilize, place the Falcon tube in a beaker filled with water and boil for 20 minutes.
- Aliquot 1mL into 1.5mL Eppendorf tubes and store at −20°C up to 12 months.
- For short term storage (<3 months), keep at +4°C.

### Synthetic Media Dropout Plates

- Prepare the following concentrated stocks in ddH20: 1.7%(w/v) Yeast Nitrogen Base (BD Difco, 233520), 5%(w/v) ammonium sulfate (Sigma, 31119), 100X amino acids and additional supplements (see below) if necessary for optimal the growth of the specific *S. cerevisiae* background.
- For the strain background used in these protocols, the following concentrated stocks need to be prepared:

- 100X Amino Acid Mix: Mix Adenine Hemisulfate (4g/L, Sigma A9126), Arginine (2g/L Sigma A5131), Aspartic Acid (5g/L, Sigma 9256), Glutamic Acid (5g/L, Sigma G1626), Isoleucine (6g/L, Sigma I2752), Phenylalanine (6g/L, Sigma P2126), Proline (4g/L, Fluka 81709), Serine (20g/L, S4500), Threonine (10g/L, T8625), Tryptophan (4g/L, T0254), Tyrosine (0.4g/L, Sigma T3754), Valine (12g/L, Fluka 94619) in ddH_2_0.
- 100 X Histidine (Sigma H8125): 2g/L in ddH_2_0.
- 100 X Leucine (Sigma L8000): 8g/L in ddH_2_0.
- 100 X Lysine (Sigma L5626): 4g/L in ddH_2_0.
- 100 X Methionine (Sigma M9625): 2g/L in ddH_2_0.
- 100 X Uracil (Sigma U0750): 2g/L in dd_2_0.
- Filter sterilize all solutions using a filter with 0.22 μm pore size.
- Store at +4°C for up to one year.
- For Synthetic Defined Media Dropout plates, add 2% (w/v) Agar (<u>Cas Number 9002-18-0</u>) to ddH_2_0.
- Autoclave the solution. Once the solution is down to 60 °C, add to a final concentration of 0.17% (w/v) Yeast Nitrogen Base, 0.5%(w/v) ammonium sulfate, 1X amino acids and additional supplements if necessary for optimal the growth of the specific. *S., cerevisiae* background. Autoclaved ddH_2_0 can be used to adjust volume. The amino acid or supplement used for selection during the transformation protocol is always omitted from the media.
- Store plates upside down, sealed in plastic bags at +4°C for up to 12 months.

### 10 X Taq/Vent Buffer

- Mix the following components with the indicated final concentrations: 500mM Tris-HCl(pH=9.0) (see above), 22.5mM MgCl2(CAS number 7786-30-3), 160mM NH4SO4(CAS number 7783-20-2).
- Store at −20°C for up to 2 years and +4°C for up to 6 months.

### 10X TBE buffer

- Dissolve the following components in dd_2_0: Boric acid (Sigma Boric Acid or equivalent, CAS no. 10043-35-3), Trizma Base (Sigma Trizma Baes or equivalent, CAS no. 77-86-1) and 0.08M EDTA solution (see above) to a final concentration of 0.9M, 0.9M and 0.01M, respectively.
- Store at room temperature for up to one year.
- If components precipitate, autoclave to dissolve.

### Electrophoresis gel

- Dissolve the required concentration (e.g. 2% (w/v)) ultrapure Agarose (Thermofisher UltraPure Agarose, catalog no. 16500500, or equivalent) in 1X TBE buffer in a 250mL Erlenmeyer flask.
- Heat in a microwave until boiling.
- Once boiling, remove from the microwave and shake to dissolve the agarose. Handle the flask with heat resistant gloves.
- Pour in an electrophoresis chamber equipped with the desired combs.

### 6X Loading Dye

- Dissolve 40% (w/v) sucrose (Sigma Sucrose or equivalent, CAS no. 57-50-1) and 0.05% (w/v) Amaranth (Sigma Amaranth or equivalent, CAS no. 915-67-3) in ddH_2_0.
- Store at +4°C for up to a year.
- Before using to load samples into an electrophoresis gel, add 0.5% (v/v) GRGreen Nucleic Acid stain (LabGene Scientific, cat. No. EG1071) and vortex until well mixed.

## DISCUSSION

### Background Information

Since the harnessing of temperature sensitive and nonsense mutations (Epstein et al., 1963), the ability of researchers to regulate expression of genes and their protein products has been immensely helpful for understanding biological function. This capability has allowed elucidation of links between genotype and phenotype (Richardson et al., 1989), gene function in temporally ordered processes (Hereford & Hartwell, 1974), and manipulation of cell and organismic physiology to aid scientific discovery of development of therapeutics or biotechnological products (Sochor et al., 2015). These discoveries were made with gene deletions or temperature sensitive mutations, which eliminate or impair protein function. Such methods, however, cannot provide dosage control and mostly allow merely qualitative observations based on complete absence of the protein product or function. Over time, natural *S. cerevisiae* promoters induced, for example, by sugars (Gal) or metabolites were used to more precisely vary gene dosage (Maya et al., 2008). More recent control schemes have included chimeric transcription factors that carry bacterial DNA binding moieties and eukaryotic transcriptional activators to induce expression of targeted genes by small molecules (Bellí et al., 1998a; McIsaac et al., 2014a; Ottoz et al., 2014a). More recently, optogenetic systems have been implemented to control gene expression in yeast (Figueroa et al., 2021). These systems typically involve two proteins that dimerize when stimulated by light (Salinas et al., 2018; Xu et al., 2018; Zhao et al., 2018). One of these partners is fused to a DNA binding domain, and the other is fused to an activation domain. Therefore, upon dimerization, they can bind a promoter with a dedicated binding site and induce transcription. Whilst these natural and artificial transcriptional controllers are widely utilized, they suffer from various drawbacks, such as persistent basal leakiness, cell-toxicity, low maximum expression capacity, high cell-to-cell variation in expression or specific media dependencies. WTC_846_ was developed to address these drawbacks (Azizoğlu et al., 2021).

WTC_846_ uses a bipartite architecture with two separate repressors to control expression of the target gene (Figure 1A). TetR fused to Tup1 represses expression of the target gene completely, especially when used in conjunction with a low translation efficiency extended Kozak sequence. This repressor is constitutively expressed, and it is known that constitutively expressed transcription factors lead to high cell-to-cell variation upon induction (Elison et al., 2017; Meurer et al., 2017; Ottoz et al., 2014). In order to reduce this variation, we use a second repressor, TetR, which also represses itself to create an Autorepression (AR) loop. We call this architecture, which combines a constitutively expressed and a negatively autoregulated repressor, Complex AutoRepression (cAR)(Azizoğlu et al., 2021). While the constitutive repressor abolishes basal leakiness of the system, the AR loop suppresses cell-to-cell variation compared to most other controllers, which use only constitutively expressed repressors/activators, and therefore exhibit high cell-to-cell variation in expression (Bellí et al., 1998b; McIsaac et al., 2014b; Ottoz et al., 2014b; Salinas et al., 2018; Xu et al., 2018; Zhao et al., 2018).

Compared to existing controllers of gene expression such as the galactose inducible system, LexA-activation domain fusions or optogenetic systems, WTC_846_ has a number of advantages. It can be turned off completely in the absence of anhydrotetracycline and can, thus, generate reversible knockout phenotypes. At full induction, expression level rivals that of the highest expressed proteins in the yeast proteome, therefore enabling overexpression studies (Azizoğlu et al., 2021; Swaffer, Marinov, et al., 2021). Additionally, WTC_846_ works in many different growth media conditions without any observable burden on growth. Finally, expression is “clamped” by the autorepression loop, so is uniform throughout the cell population. In other words, population mean is representative of single cell behavior. While there are controllers that do fulfill one or more of these criteria, WTC_846_ is unique in achieving all of them.

The protocols described here explain how to generate strains in which WTC_846_ regulates the expression level of the target genes, and provide a useful example of such control, namely, to turn off and then turn on expression of a targeted cell cycle regulatory gene, *CDC20,* in order to synchronize the culture. Such cycle synchronization is helpful when studying gene expression or other cell functions in different cell cycle phases. The better synchronized a batch culture is, the more enriched it will be for cells in the relevant cell cycle stage. This then makes it easier to get reliable readouts of, for example, cell-cycle dependent protein phosphorylation levels or protein-protein interaction information from batch cultures (Dai et al., 2018; Ewald et al., 2016).

Furthermore, we described an optimized LiAc-based transformation protocol (Basic Protocol 1) and an alternative CRISPR-Cas9 based transformation protocol (Alternative Protocol 1) for incorporating elements of the system into the genome. The transformation methods in these protocols were extensively optimized to result in high transformation efficiencies.

Depending on the controlled gene, the WTC_846_ system can be used to control diverse aspects of cell function, from metabolic reactions to properties like cell size and growth rate (Azizoğlu et al., 2021). The precision with which protein dosage can be controlled is likely to reveal and allow manipulation of dosage dependent phenotypes. For example, the system has been used recently to express the cell size control protein Whi5, where the authors required cell-cycle independent expression of Whi5 whilst keeping the set-point and population variance of cell size same as a wild type population (Swaffer, Kim, et al., 2021). This was only possible due to the low cell-to-cell variation in expression of WTC_846_ strains. Another use of the system has been to place multiple genes under identical regulation. This was achieved in a recent study, where the authors placed all 12 subunits of the RNA polymerase II under the control of WTC_846_ for overexpression (Swaffer, Marinov, et al., 2021). Furthermore, genome-wide or large-scale collections of WTC_846_ are likely to prove valuable when investigating dosage dependent epistasis interactions between genes(Costanzo et al., 2016a). Additionally, current arrest and release methods use either chemicals, temperature shifts, or inducible systems that cannot be fully shut down (Costanzo et al., 2016b; Ewald et al., 2016). Therefore, when controlling cell cycle genes to synchronize or regulate cell cycle state, WTC_846_ represents a better, more penetrant and less physiologically perturbing alternative to current arrest and release methods.

### Critical Parameters

#### Achieving high transformation efficiency

The two main factors that will affect the success of all transformation protocols described here are the quality of the cell culture and the quality of the DNA to be used in the transformations. Cells should always be in exponential phase during transformation, having been cultured in fresh rich media for at least 3 hours or 2 cell divisions prior to beginning (Ito et al., 1983). Performing the protocol with too few (<10 million/mL) or too many (>30 million/mL) cells is likely to reduce the number of correct transformants. In our hands, plasmid or PCR product DNA that was recently prepared and resuspended in double distilled water gave the best results. Increasing the amount of DNA used for transformation can increase the number of correct transformants (Ito et al., 1983).

#### Achieving high efficiency with the CRISPR-based transformation

All points discussed above also apply to CRISPR-based transformations within Alternative Protocol 1. Additionally, the choice of cut site (defined in Alternative Protocol Step 4) has been reported by users to influence efficiency. There are two homologous regions between the repair template and the genome, designed in Alternative Protocol Step 1 or 2. For maximum efficiency, the cut site should be as close as possible to one of these homologies if possible (i.e. at most 20-30 bp distance between the cut site and the start of the homology region).

It is crucial to see a distinct growth rate difference between +/− β-estradiol conditions during the spotting assay before continuing with the protocol. Lack of a distinct growth rate difference will usually indicate a CRISPR plasmid that assembled without sgRNA sequences, and such a CRISPR Mid-strain should be discarded.

#### Determining optimum expression level for WTC_846_ strains

Once a *WTC_846_::GOI* strain is isolated and established, it should be maintained with an anhydrotetracycline concentration in both liquid and solid medium that results in near endogenous level expression of the controlled gene. This is especially important for essential genes, since an increase or a decrease in their protein levels can alter growth rates as well as have other side effects. These could range from DNA damage to cell cycle arrest depending on the function of the controlled gene. See Strategic section on how to estimate the correct aTc concentration for different controlled genes. It is important to note that on solid culture, the required inducer concentration is higher, likely due to slow diffusion of aTc into the growing colony and depletion of aTc from the immediately adjacent solid medium. For most endogenous genes, information on endogenous expression levels is available from the Saccharomyces Genome Database (www.yeastgenome.org).

#### Non-steady state use of WTC_846_ to shut off and induce gene expression

Induction of gene expression with anhydrotetracycline starts within 30 minutes, and depending on the degradation rate of the protein product, reaches steady state within, at most, around 7 hours (Figure 5 and 11). Time to reach steady state depends on the protein; the more stable the protein, the more time after induction it will take to reach steady state expression level.

**Figure 11.**
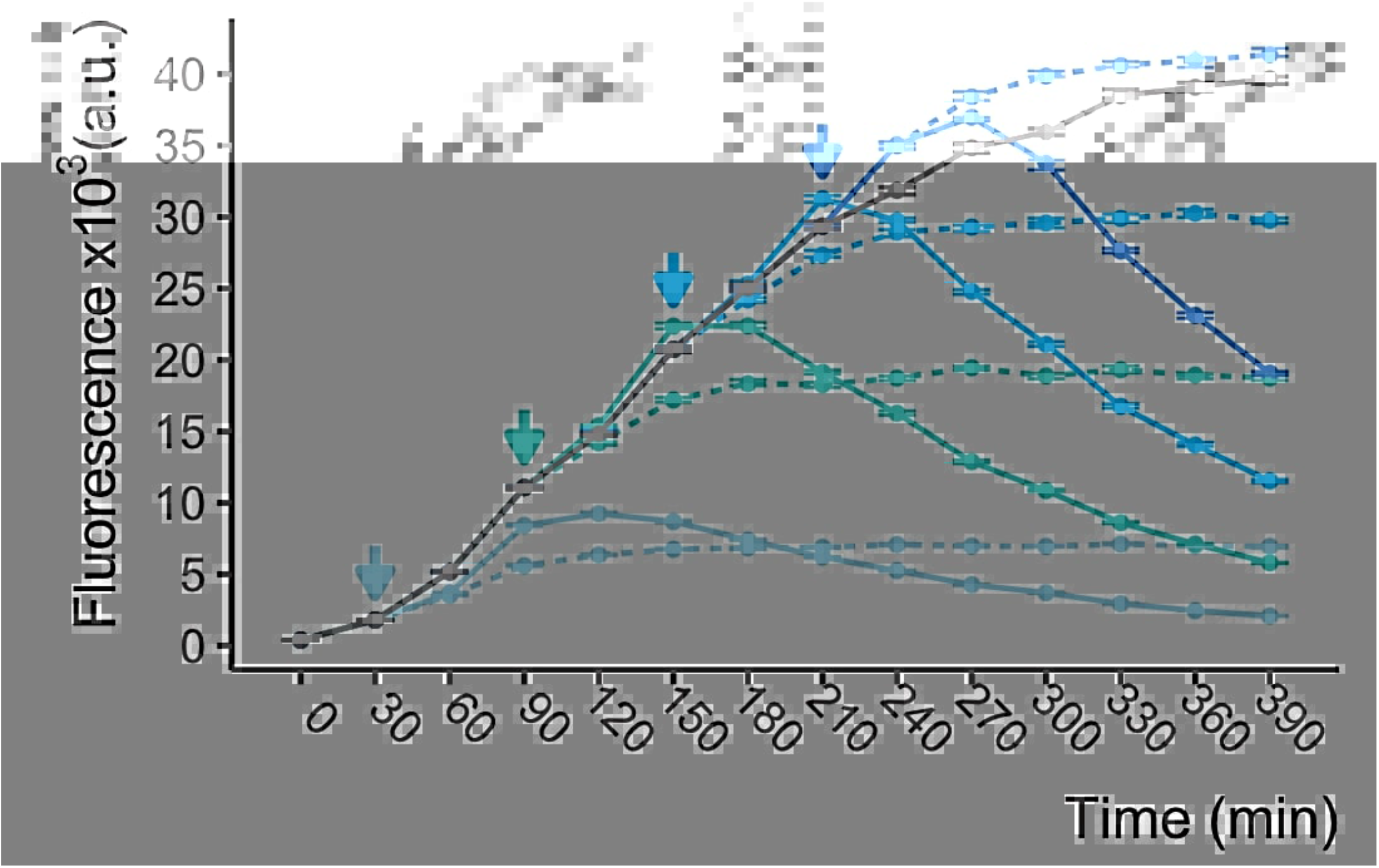
Induction and shutoff kinetics in the *WTC_846_::Citrine* strain. Cells were in early exponential phase in liquid YPD when 600ng/mL aTc was added to the culture. The main culture (grey) was allowed to fully induce over a period of 390 minutes in the presence of aTc. At times indicated by the colored arrows, samples were taken from the main culture, washed, and resuspended in either YPD without aTc (colored solid lines) or YPD without aTc but with 70μg/mL of translation inhibitor cycloheximide (colored dashed lines) to start shutoff. Cycloheximide (CHX) inhibits further protein production and is used to get an indication of residual folding and subsequent fluorescence emission of the remaining, newly translated Citrine protein. Since CHX also inhibits cell division, dilution of the Citrine fluorescence is not observed in these samples. Citrine fluorescence was measured with flow cytometry (488nm excitation, 530/30nm emission). Circles represent the median of the population and the error bars indicated 95% confidence interval. Figure was reproduced with permission from (Azizoğlu et al., 2021).

Shutoff of targeted gene expression requires complete removal of the inducer from the media, after which transcription of the gene will cease almost immediately (Figure 11). Regardless of whether the cells come from liquid or solid medium, careful wash steps should be performed with liquid medium without inducer. If the starting inducer concentration was high, the concentration of the targeted protein in the cells will also be high. Therefore, especially if the protein product has a low degradation rate, it may be necessary to dilute the cells into fresh medium without inducer and continue growing them until the cellular level of the starting protein drops to zero. In this case, collect, wash, and dilute a portion of the culture into fresh medium without inducer. Given that time to zero will depend on the initial inducer concentration, stability of the protein product and the thoroughness of the wash steps, a successful shutoff experiment will require trials to establish a robust protocol for each different gene being controlled.

### Troubleshooting

See Table 1 for a list of common problems with the protocols, their causes, and potential solutions.

### Understanding Results

We provide data collected using a *WTC_846_::Citrine* strain as examples of a typical liquid culture dose response (Figure 5), solid media dose response (Figure 6), and liquid media induction and shutoff kinetics (Figure 11) of the WTC_846_ system. Here, to guide use of newly constructed strains, we discuss how to extrapolate from the behavior of the well-studied *WTC_846_::Citrine* to predicting how different WTC_846_ strains might behave regarding dose response, and induction and shutoff kinetics, and how the behavior of the strains will vary depending on the media, temperature, and other variables.

#### Dose response

In WTC_846_, the targeted gene is controlled by the promoter P_7tet.1_ (Figure 1A). This promoter is based on the endogenous P_TDH3_ promoter of *S. cerevisiae* and has a similar maximum expression level as assessed by fluorescence of strains where both promoters were driving Citrine expression(Azizoğlu et al., 2021). Therefore, the maximum protein concentration reached by fully inducing WTC_846_ with 600 ng/mL aTc is roughly equivalent to *TDH3* protein concentration inside the cell. Since the median TDH3 molecules per cell is estimated to be around 2 million in YPD (Ho et al., 2018), appropriate aTc concentrations for desired expression levels of other genes can be calculated, provided their median expression level is known and the degradation kinetics between TDH3 and that protein are similar. This information is available for most genes on the Saccharomyces Genome Database (www.yeastgenome.org) (Cherry et al., 2012). A more detailed explanation of this calculation is given in Strategic Planning.

#### Induction and shutoff

Induction of the *WTC_846_::Citrine* strain starts almost immediately and Citrine fluorescence is already detectable at 30 minutes (Figures 5 and 11). For a stable protein like Citrine, reaching steady state takes around 6 to 7 hours. Time to steady state will, however, depend on two factors: stability of the GOI and the growth rate in the specific media used for the experiment. The higher the stability of the GOI protein product (i.e. the lower the degradation rate), the longer it will take to reach steady state. Similarly, since a slower growth rate reduces the dilution rate of the protein, the slower the growth rate, the longer it will take to reach steady state.

In terms of shutoff, Figure 11 demonstrates that transcription of the GOI ceases almost immediately, yet complete depletion of protein takes a long time. This is also apparent in the arrest and release assay depicted in Figure 10. In the case of Citrine, protein degradation is negligible, and fluorescence is diluted only with cell division. Additionally, the higher the starting Citrine concentration, the longer the shutoff takes. Therefore, shutoff kinetics when using WTC_846_ with any controlled gene will depend on the degradation kinetics of the specific protein product and the initial starting concentration of the protein. Furthermore, the specific media conditions and temperature of incubation will affect the growth rate, therefore affecting shutoff kinetics. The slower the growth rate, the longer it will take to dilute all existing protein through cell division and to reach shutoff.

## Considerations

Basic Protocol 1 and 2 both take 3 days, with a total of 1-2 hours of bench time. Alternate Protocol 1 takes 7-8 days, with 3-4 hours of bench time. All three protocols require preparation steps starting from 3 days in advance. Basic Protocol 3 takes 1.5 days, with 1 hour of bench time.

## DATA AVAILABILITY STATEMENT

Most of the data presented here is openly available in a public repository that issues datasets with DOIs (https://doi.org/10.3929/ethz-b-000488967). The rest is available on request from the authors.

## ACKNOWLEDGEMENTS

This work was supported by the Swiss National Science Foundation as part of the Molecular Systems Engineering NCCR and by grant R21CA223901 from the NCI to RB.

